# Curvature of the retroviral capsid assembly is modulated by a molecular switch

**DOI:** 10.1101/2020.12.26.424437

**Authors:** Tyrone Thames, Alexander J Bryer, Xin Qiao, Jaekyun Jeon, Ryan Weed, Kaylie Janicki, Bingwen Hu, Peter L. Gor’kov, Ivan Hung, Zhehong Gan, Juan R Perilla, Bo Chen

**Author notes:** These authors contributed equally to this work (Tyrone Thames, Alexander J Bryer, Xin Qiao).

## Abstract

During the maturation step, the capsid proteins (CAs) of a retrovirus assemble into polymorphic capsids, whose acute curvature is largely determined by insertion of 12 pentamers into the hexameric lattice. Despite years of intensive research, it remains elusive how the CA switches its conformation between the quasi-equivalent pentameric and hexameric assemblies to generate the acute curvature in the capsid. Here we report the high-resolution structural model of the RSV CA T=1 capsid. By comparing with our prior model of the RSV CA tubular assembly consisting entirely of hexameric lattices, we identify that a dozen of residues are the key to dictate the incorporation of acute curvatures in the capsid assembly. They undergo large torsion angle changes, which result a 34° rotation of the C-terminal domain relative to its N-terminal domain around the flexible interdomain linker, without substantial changes of either the conformation of individual domains or the assembly contact interfaces.

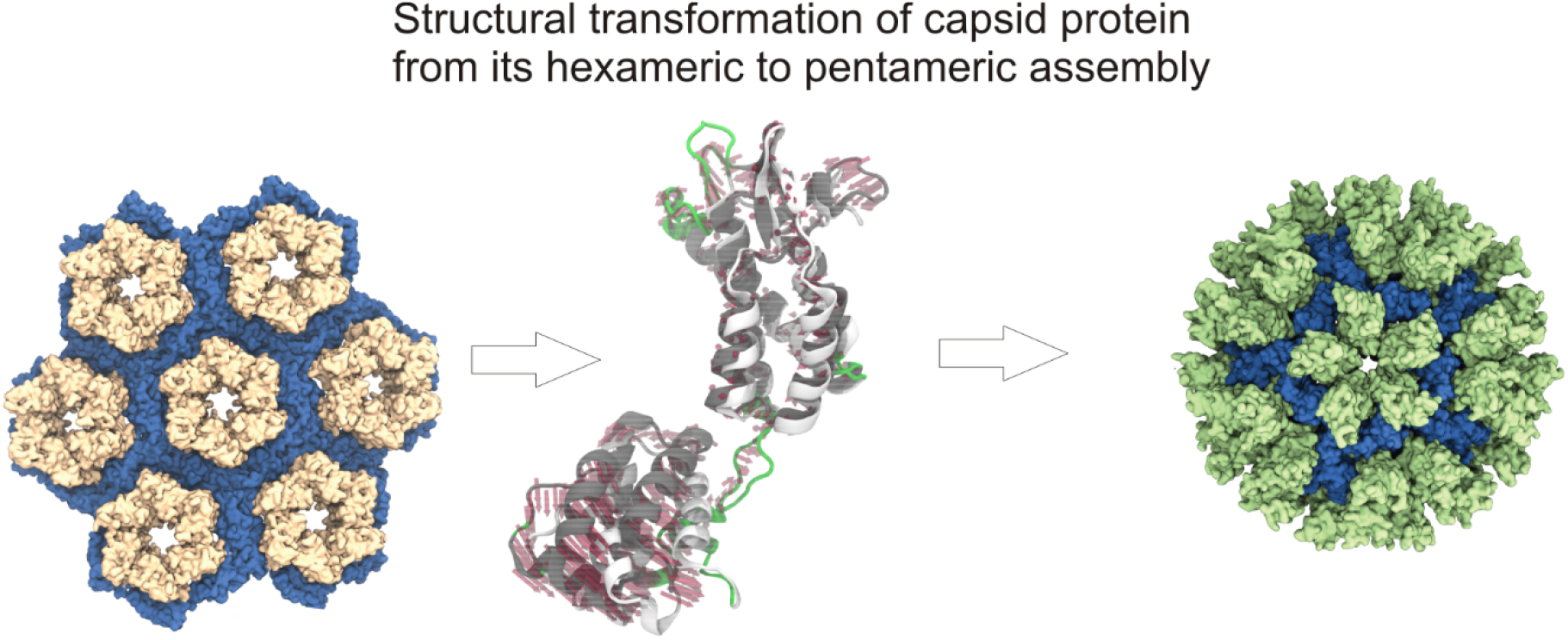

## Introduction

A mature retroviral capsid is assembled by more than 1000 copies of the capsid protein (CA) during a process called maturation in its life cycle to finalize its capsid assembly.*^1^* Despite limited sequence similarity, monomeric orthoretroviral CAs share nearly identical 3D folding.*^2^* Briefly, a CA contains two independently folded domains joint by a short flexible linker,*^3^* with the N-terminal domain (NTD) comprising a short β-hairpin followed by seven α-helices,*^4, 5^* and the C-terminal domain (CTD) consisting of a short 3__10__ helix followed by four additional α-helices.*^6^* Intriguingly, these CAs form polymorphic capsids of distinct morphologies for different viruses, including the tubular, polyhedral and conical structures.*^2, 7^* It was shown that the correct capsid assembly is critical to viral propagation and infectivity.*^1, 8^* Hence, CAs and their assemblies are promising antiviral drug targets to overcome the fast mutation hurdle.*^9, 10^* However, due to the strong polymorphism,*^2, 11^* the detailed molecular basis of retroviral capsid assembly remains elusive.

A key unresolved question is the mechanism of the curvature management in the retroviral capsid assembly. Based on analyses of *in vivo* and *in vitro* assemblies, the polymorphism and peculiar shapes of retroviral capsids are assembled by insertion of 12 CA pentamers at varying locations in their hexameric lattice.*^7, 12^* A great deal were learned about the hexameric assembly of retroviral CAs by studying various *in vitro* systems, while the pentameric assembly as the cause of acute assembly curvature remains largely elusive.*^11^* Ganser *et al.* obtained the first high resolution structural model of the hexameric assembly formed by HIV CA R18L mutant, and identified three assembly interfaces unambiguously for the first time:*^13^* the first three helices form the NTD-NTD interface that bind adjacent subunits into individual hexamers, while the CTDs are stabilized by the NTD-CTD interface between helix 4 and 8 within each subunit. Adjacent hexamers are bound together by the CTD dimer interface across helix 9 into a complete hexameric lattice. Subsequently, Pornillos *et al.* obtained crystal structures of isolated HIV CA hexamers*^14, 15^* and isolated pentamers*^16^* by crosslinking of mutant HIV CA and Ccmk4 fused CA. Both pentamer and hexamers exhibit interfaces similar to the cryogenic electron microscopy (cryoEM) model.*^13^* In addition, they found that NTD-NTD and NTD-CTD interfaces are solvated with water, with little variations across all hexamer models. Instead, CTDs at the edge of isolated hexamers adopt a variety of orientations. This was interpreted as the cause of assembly plasticity. The charged residues closer at NTD-NTD interface in the CA pentamer were suggested to act as the switch of assembly. However, these crosslinked oligomers are in isolated state with disrupted CTD dimer interface. Around the same time, Cardone *et al.* and Hyun *et al.* obtained ~ nm resolution models of T=1 and T=3 Rous sarcoma virus (RSV) CA capsids, showing interfaces similar to those in HIV CA hexamers. Their results proved that different retroviruses adopt a common capsid assembly structure and potentially similar curvature control mechanism.*^17-19^* However, the limited resolution does not allow distinction of CA pentamers and hexamers at the molecular level. Subsequently, Zhao *et al.* obtained an 8 Å resolution model of the HIV CA tubular assembly by cryoEM. In this curved hexameric assembly, a fourth interface, the CTD trimer interface, was discovered, between helix 10 and 11,*^20^* critical to the tubular curvature. Atomic resolution models of different shaped mature HIV capsids were constructed,*^20^* using molecular dynamics (MD) simulation to insert isolated pentamers*^16^* into their tubular model. The models indicate that variations of CTD dimers are needed for different curvatures around the capsid, which agrees with earlier work by Pornillos et al.*^16^* In the 8.8 Å resolution cryoEM model of the authentic HIV capsid,*^21^* pentamers seem to adopt different NTD-NTD and NTD-CTD interfaces than those in crosslinked hexamer and pentamers.*^14, 16^* However, details could not be resolved at the limited resolution. Meanwhile, analysis of different RSV CA planar hexameric sheets at 4 Å resolution by Bailey *et al.* indicates that both NTD-NTD and CTD dimer interfaces are rigid, but the NTD-CTD orientation varies around the NTD-CTD interface in different sheets.*^22^* However, no curvature is present in these assemblies. Recent atomic resolution X-ray diffraction models of native HIV CA hexamer crystals identified prevalent hydrogen-bonds (H-bonds) at all interfaces.*^23^* They also showed that variation of the hydration level triggers changes of NTD-CTD orientations around the interdomain linker, which attributes to the assembly plasticity. In addition, they found notable differences at the CTD dimer interface from that in isolated hexamers of crosslinked mutant HIV CA,*^14, 15^* with a tighter CTD trimer interface.*^20^* Meanwhile, solid state NMR (ssNMR) were applied to study various HIV CA assemblies.*^24–31^* Residues at the ends of each domain and in the cyclophilin A binding loop were shown to be highly mobile, while the linker between two domains undergoes millisecond motions,*^25, 26^* which could alter the interdomain orientation and cause assembly polymorphism. However, no major distinction at the molecular level could be identified between CAs in the pentameric and hexameric assembly.*^25, 27, 28^* Hence, it is still not clear how retroviral CAs switch between quasi-equivalent pentamers and hexamers, and controls assembly curvature.

In our previous work, we obtained the first high resolution model for the RSV CA tubular assembly by combining ssNMR and cryoEM constraints.*^32^* We showed that the curvature variations in the hexameric lattice is mediated by varying side chain configurations, while the backbone structure remains largely unchanged. In this work, we report the first high resolution structural model of the RSV CA in its T=1 capsid assembly, with 1.97 Å root-mean-square deviation (RMSD) of Carbon Cα among CA monomers. Molecular dynamics flexible fitting (MDFF) allows site-specific resolution of the molecular details of CA pentamers by combining secondary structural constraints from our ssNMR results with prior published cryoEM model (EMDB 1862) at 8.5 Å.*^19^* The model reveals a simple mechanism of the RSV CA to switch between the hexameric and pentameric assembly: the subunit undergoes a 34° interdomain rotation via the flexible interdomain linker, which causes large torsion angle changes to only about a dozen residues, with little adjustment of domain structure or interface contacts.

## Results and Discussion

Wild-type RSV CAs tend to form either tubes or large polymorphic structures comprising mostly hexamers.*^33^* To prepare assembly comprising pentamers, we adopted the I190V RSV mutant.*^34^* Its solution at 6 mg/ml forms predominantly small spherical structures mixing with equal volume of 1 M sodium phosphate buffer at pH 8. To assess the morphology of the assemblies, negatively stained transmission electron microscopy (TEM) images were recorded. A representative view is shown in Figure 1A. The statistics of the assembly diameter was analyzed. The histogram is shown in Figure 1B. As shown, the average size of the spherical assemblies is about 18.5 nm, consistent with the expected value of T=1 capsid.*^17^* Hence, it confirms that our assembly sample consists nearly entirely of CA pentamers.

**Figure 1.**
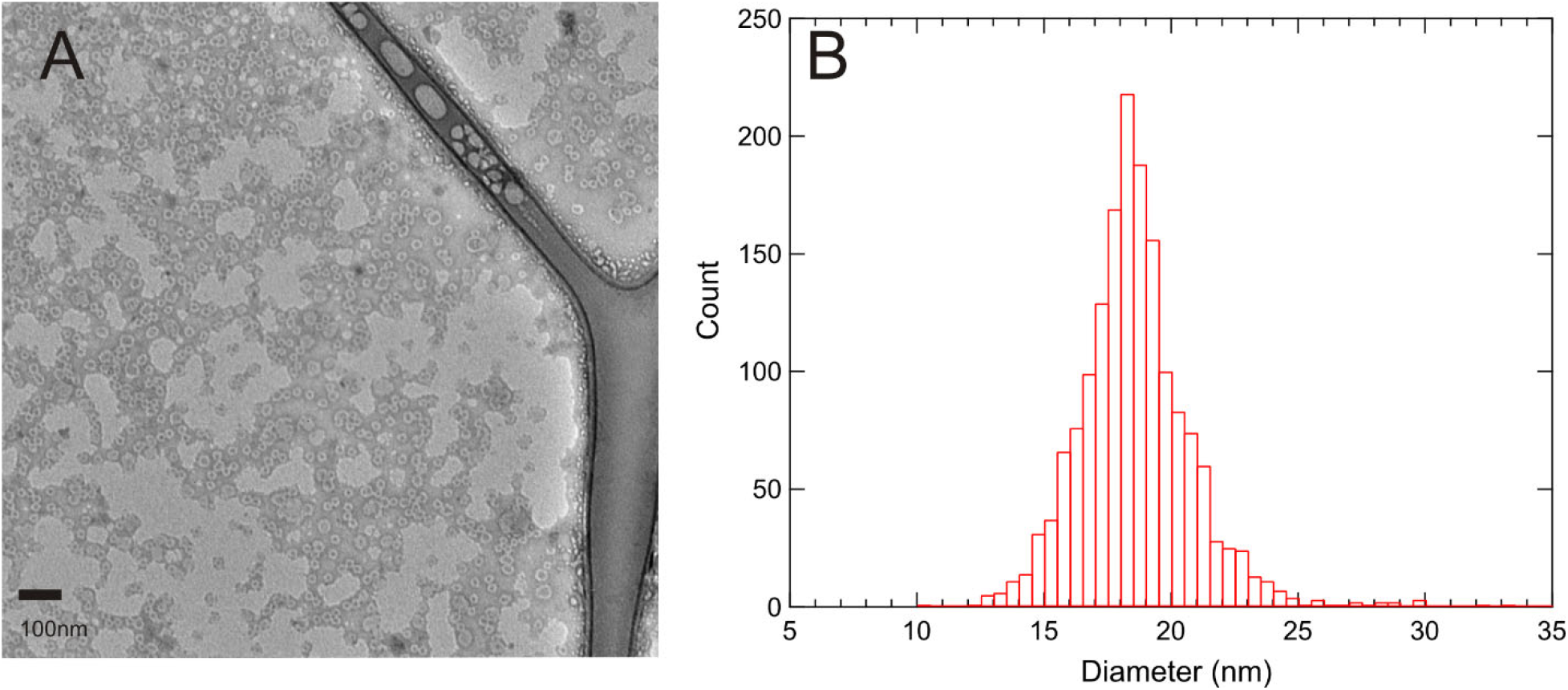
(A). Negatively stained TEM image of the RSV CA I190V T=1 capsid assembly. The operating magnification is 15,000-fold. (B). The average diameter of observed assemblies is ~18.5 nm as shown by histogram of the distribution of assembly diameters.

To perform NMR experiments, uniform ^13^C, ^15^N labeled CA (U-CA) was expressed. Its assembly produces 2D spectra of excellent resolution. A 2D ^15^N-^13^C correlation spectrum with cross peaks created by 50 ms dipolar assisted rotational resonance*^35^* is shown in Figure 2A. The typical linewidth is about 0.7 ppm along the ^13^C dimension and 1.2 ppm along the ^15^N dimension, comparable to that of our prior RSV CA tubular assembly under moderate magic angle spinning (MAS),*^32^* and similar to that of other highly ordered protein structures*^36–39^* and HIV CA assemblies.*^24–31^* It indicates that RSV CA adopts highly order structure at the molecular level, consistently with our TEM evaluation.

**Figure 2.**
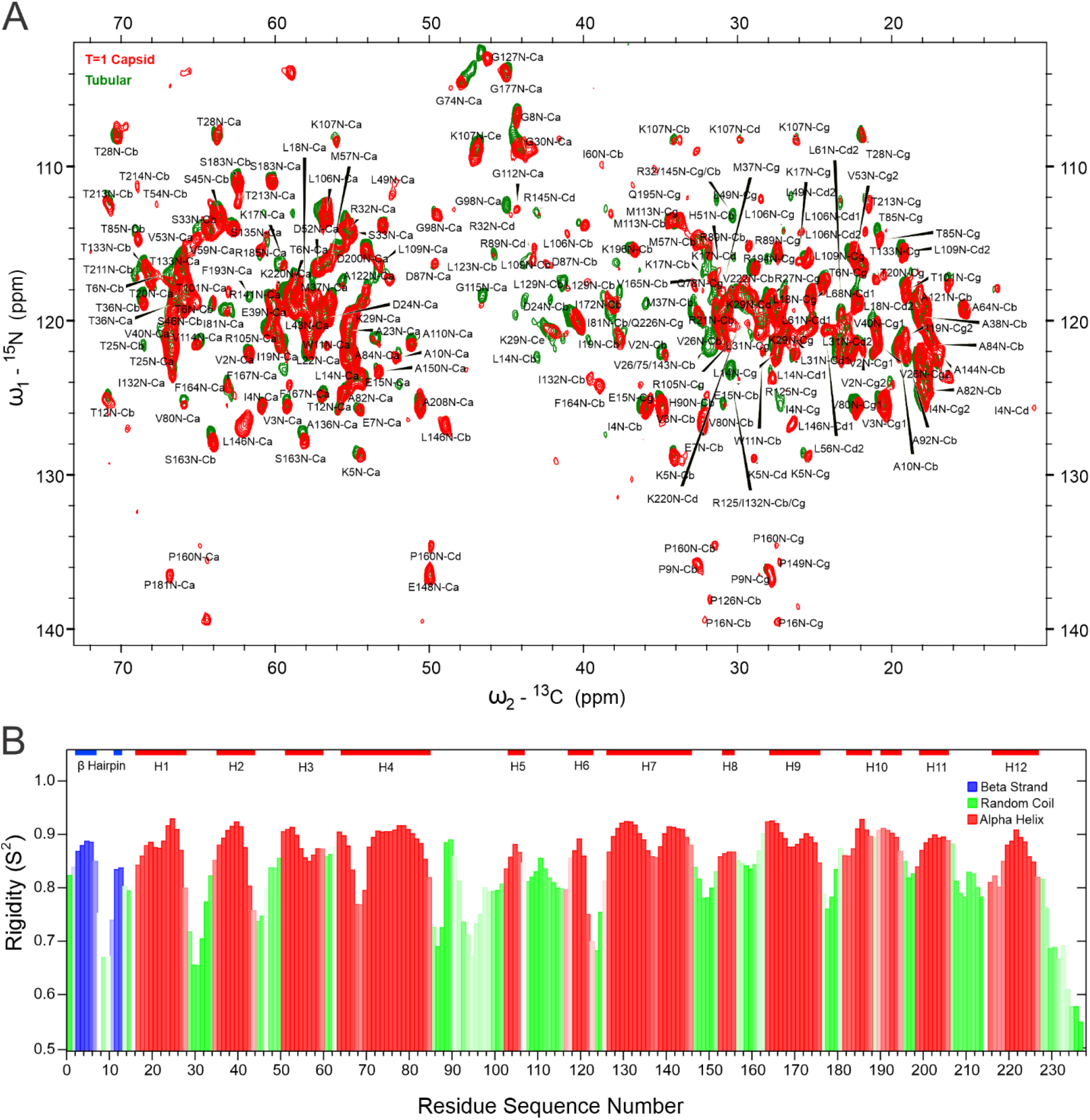
Dynamics and secondary structure of the T=1 capsid assembly by the RSV I190V CA derived from ssNMR assignments. (A). 2D ^15^N-^13^C correlation spectra overlay of the T=1 capsid assembly by the RSV I190V CA (red) and the tubular assembly by the wild-type RSV CA (green). (B). Site-specific prediction of secondary structure and rigidity by TALOS-N^41^ from chemical shift assignments. Bar height represents the S^2^ order parameter, a measurement of rigidity. Bar saturation represents prediction confidence. The thick color lines at the top highlight segments of secondary structured units, blue for β-strands and red for α-helices.

To perform sequential assignments, a series of 2D and 3D correlation spectra were recorded with the U-CA, as detailed in Table S1 in the supporting information. All congested resonances in 2D spectra can be resolved in the 3D NCaCX and partially resolved in 3D NCOCX spectra. We note that only one set of resonances were identified for each residue, which suggests that the RSV CA adopts a unique and well-defined conformation in the T=1 assembly. Manual sequential assignments were performed by backbone walk to link resonances in 3D NCaCX and NCOCX. Together, 115 residues were sequentially assigned. A stretch of the sequential walk assignments between residues 53–65 is shown in Figure S1 in the supporting information.

Additionally, 104 well resolved residues were identified in either 3D NCaCX or NCOCX spectra. Although backbone walk cannot be achieved for these residues, they exhibit nearly identical signals as in our prior tubular assembly. Hence, there is little uncertainty about their identities.

Resonances in these 3D correlation spectra were acquired by cross polarization transfer, mediated by anisotropic dipolar interactions (through space). Hence, only residues in rigid segments in the assembly would contribute. To detect residues in the flexible segments, INEPT-TOBSY spectrum were acquired with through bond polarization transfer,*^40^* shown in Figure S2 in the supporting information. Five residues were identified with narrow linewidth comparable to solution NMR and chemical shifts close to random coil values. These residues undergo fast motion like in the soluble state. They are assigned to P230, L231, T232, G235, and I236, as those at the tail of the CTD revolved in our prior tubular assembly work.*^32^* We also note that signals in the INEPT-TOBSY spectrum of our T=1 assembly sample are much weaker than those of the tubular assembly. It implies that these residues at the CTD tail are probably not as mobile as those in the tubular assembly, thus attenuated the signal intensity.

Altogether, 224 residues were assigned for the 237-residue RSV CA in its T=1 capsid assembly. The assignments were deposited in Biological Magnetic Resonance Data Bank (BMRB), with BMRB ID 50550. They are also shown in Table S2 in the supporting information. The following residues are missing from both through bond and through space transferred spectra: residues 47, 50, 88, 95, 117, 157, 173, 218, 219, 229, 233, 234, and 237. Except for residue 173 in the middle of helix 9, the missing residues in our T=1 capsid assembly are all in the flexible loop regions between helices. It is most likely because their motions are in the range not sensitive for ssNMR detection. In contrast, missing residues in the tubular assembly were residues 150, 229, and 230.*^32^* In addition, we note the average signal intensity in our T=1 capsid assembly is nearly 2.5 fold weaker than that in the tubular assembly,*^32^* while the spectral linewidth and acquisition conditions (experiment setup and the amount of proteins in NMR rotors) are comparable. This suggests that the molecular ordering in the hexameric and pentameric assembly is comparable, and the weaker signal intensity in the pentameric assembly is due to the shift of the overall protein dynamics towards time scale insensitive to ssNMR detection.

Based on the assignments, the secondary structure composition and local dynamics were derived with TALOS-N,*^41^* shown in Figure 2B. Overall, the RSV CA in the T=1 capsid assembly adopts a folding nearly identical to that in the tubular assembly,*^32^* with the exception for a small number of residues. This notable difference can be better appreciated by overlay of the 2D ^15^N-^13^C correlation spectra, as shown in Figure 2A: the RSV CA demonstrates similar resonances in either T=1 or tubular assembly, except for about a dozen residues exhibiting shifted resonances in both Cα and side chain regions. This is in clear contrast to the comparison of tubular assemblies with a twofold change of diameters (average diameter ~ 60 nm vs. 130 nm), which showed nearly identical Cα resonances for all residues, with shifted signals associated with side chains.*^32^* It proves that CAs have to reconfigure both their backbone and side chains to adapt to the pentameric assembly, while curvature changes in the hexameric lattice can be accommodated by side chain rearrangements alone.

To obtain the structural model for the RSV CA T=1 capsid, data-guided simulations were performed to combine our ssNMR constraints with the 8.5 Å cryoEM density (accession code EMDB 5772),*^19^* with protocols described in supporting information. The simulated model described 60 CA monomers, arranged in a T=1 icosahedron, solvated with TIP3P water*^42^* and ionized to a salt concentration of 500 mM NaCl (Figure S4A and S4B). Figure S4C shows RSV CA monomers extracted from a system of the solvated T=1 capsid simulation and aligned by the backbone atoms, with 1.97 Å RMSD of Cα among CA monomers. This alignment shows the highly uniform structure of subunits in the T=1 capsid assembly, similar to those in the tubular model. Therefore, our model allows analyses to pinpoint the distinct RSV CA conformation at a site-specific resolution in different assemblies.

Next, we computed the torsion angles of the RSV CAs measured directly from the *in silico* tubular and T=1 assembly models. Their differences are plotted in Figure 3B. The results agree remarkably well with those predicted by TALOS-N*^41^* derived directly from NMR chemical shifts (Figure 3A).

**Figure 3.**
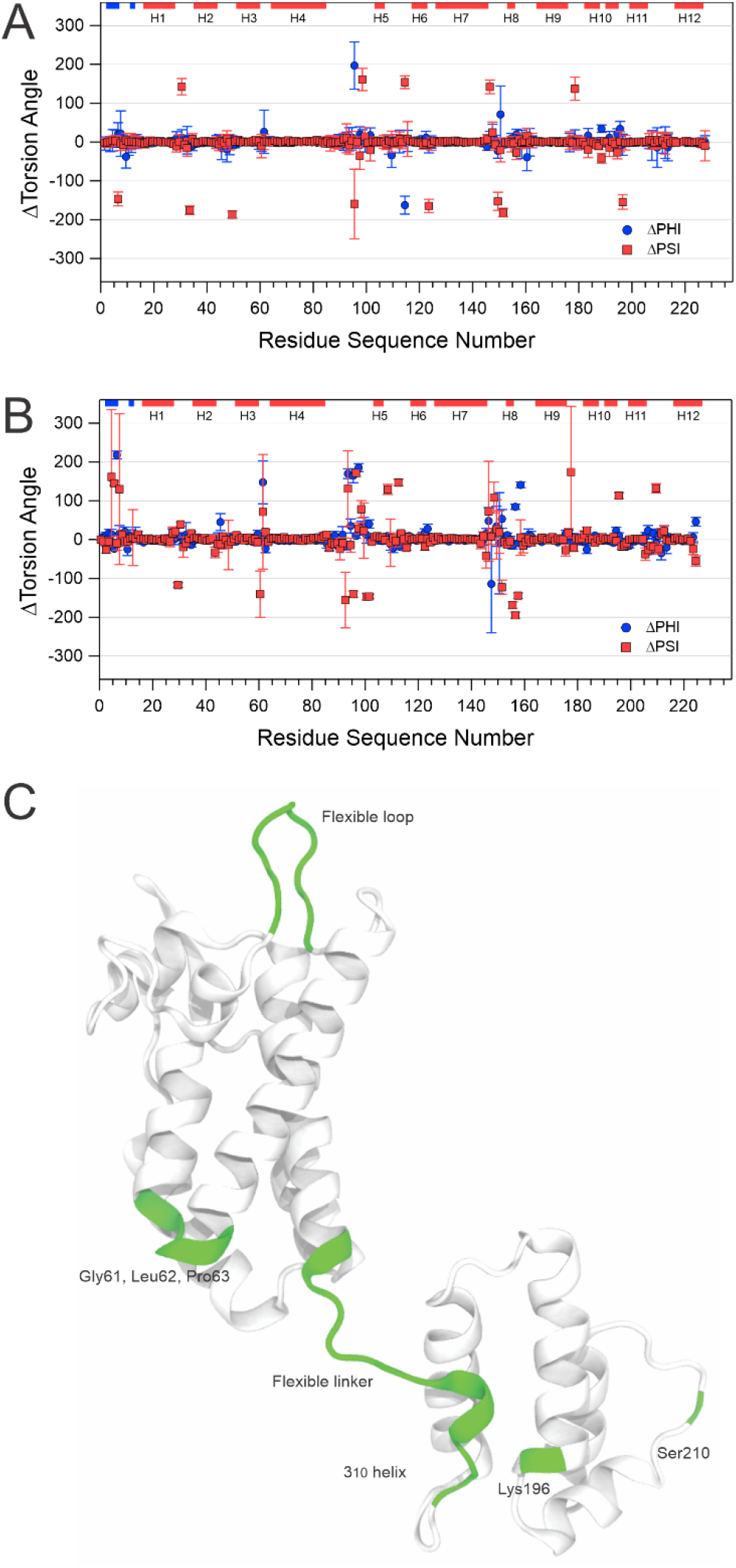
Quasi-equivalent conformation of the RSV CA in its T=1 capsid and tubular assembly. (A) and (B) are the torsion angle differences derived by TALOS-N^41^ from NMR assignments, and computed *in silico* from the T=1 capsid and prior tubular model.^32^ The thick color lines at the top highlight segments of secondary structured units, blue for β-strands and red for α-helices. (C). Residues with large torsion angle differences between two assemblies colored in green on a subunit from the T=1 capsid model.

Consistently, the regions in the RSV CA with the largest torsion angle differences are located in the β-hairpin around residues 1-16, the flexible loop and flexible linker regions encompassing residues 85-100 and 145-156, respectively (Figure 3C). Particularly, torsion angle differences based on our NMR assignments pinpoint a dozen residues with torsion angle differences greater than 100°. Their assignments in the 3D NCaCX spectra are plotted in Figure S3 in the supporting information. They reveal the site-specific changes of the RSV CA to switch from quasi-equivalent hexameric to pentameric assembly to induce acute curvature in assemblies.

Our T=1 assembly model also resolves the four intermolecular CA interfaces, critical for capsid stability and morphology,^31^ shown in Figure 4. Interestingly, torsion angle analysis shows that differences in backbone dihedral angles of these interfacial residues are limited compared to other regions of CA, such as residues of the flexible linker and other flexible regions. In the NTD, residues Leu61 and Gly62 situated near the H1-H2-H3 bundle show notable torsion angle differences although they are positioned away from the helical bundle interior (Figure 4C). Residues comprising the NTD-CTD interface (Figure 4E) show consistent torsion angles, while Ser178 – downstream of H8 and the residues which interface with H4 and H11 – shows dramatic change of the psi dihedral angle. Ser178 precedes the dimeric interface in CA which, like other interfaces, does not demonstrate large torsion angle differences among assembly models. This observation follows the trend of torsion angle differences concentrated among residues preceding and succeeding structured interfaces. Continuing down the CA sequence and preceding the trimer interface is Lys196, which additionally substantiates our observation of where torsion angles differ in distinct CA assemblies.

**Figure 4.**
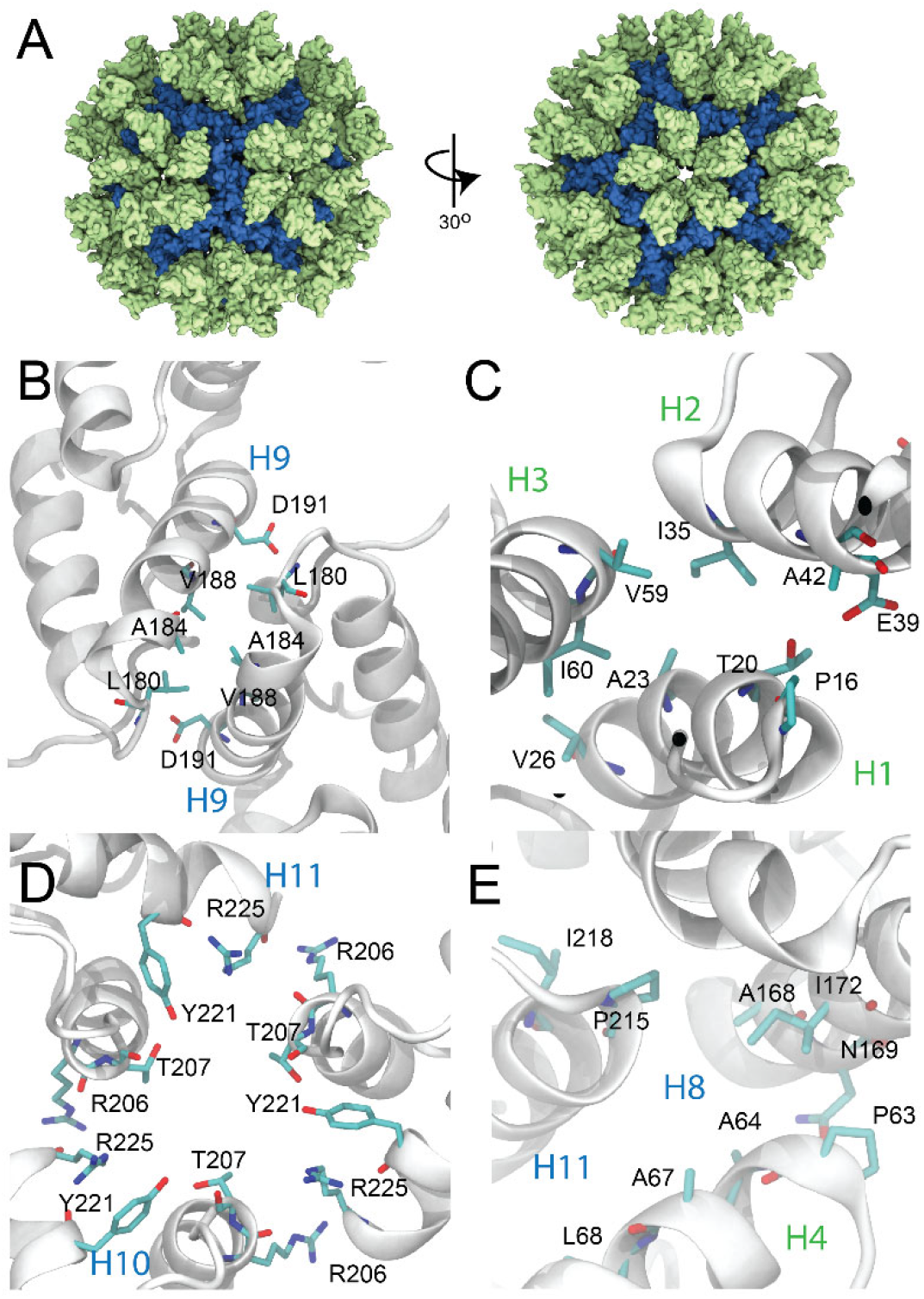
The four intermolecular assembly interfaces in the T=1 capsid model in (A) at the residue specific level. (B). The dimeric interface is between helix 9 of adjacent subunits. (C). The NTD-NTD interface comprises helix 1 to 3 of each subunit in the pentamer. (D). The CTD trimeric interface involves helix 11 and 10. (E). The NTD-CTD interface is between helix 8 and 11.

As shown in Figure 4A to D, close inspection confirms that the same set of residues are employed to stabilize each interface in both pentamers in the RSV CA T=1 capsid assembly, and hexamers in the tubular assembly.*^32^* Thus, the molecular differences in torsion angles do not cause large variations of assembly interfaces in the CA pentameric assembly compared to its hexameric assembly. It shows the remarkable efficiency of the viral CA to achieve assembly versatility with minimum modulation.

Therefore, our above analyses show only subtle differences exist in the RSV CA folding at the molecular level, in hexameric vs. pentameric assemblies. Then, the question arises, how do such subtle differences in folding translate into one less subunit in the quasi-equivalent assemblies?

To answer this, we first aligned the monomers extracted from the RSV tubular and T=1 assembly by their backbone atoms in the NTD, as shown in Figure 5A. Then, Principal Component Analysis (PCA) were applied to identify the dynamical mode or essential component that set apart the molecular folding in the hexameric vs. pentameric assembly, with their contribution ranked by the overall variance of the dataset.

**Figure 5.**
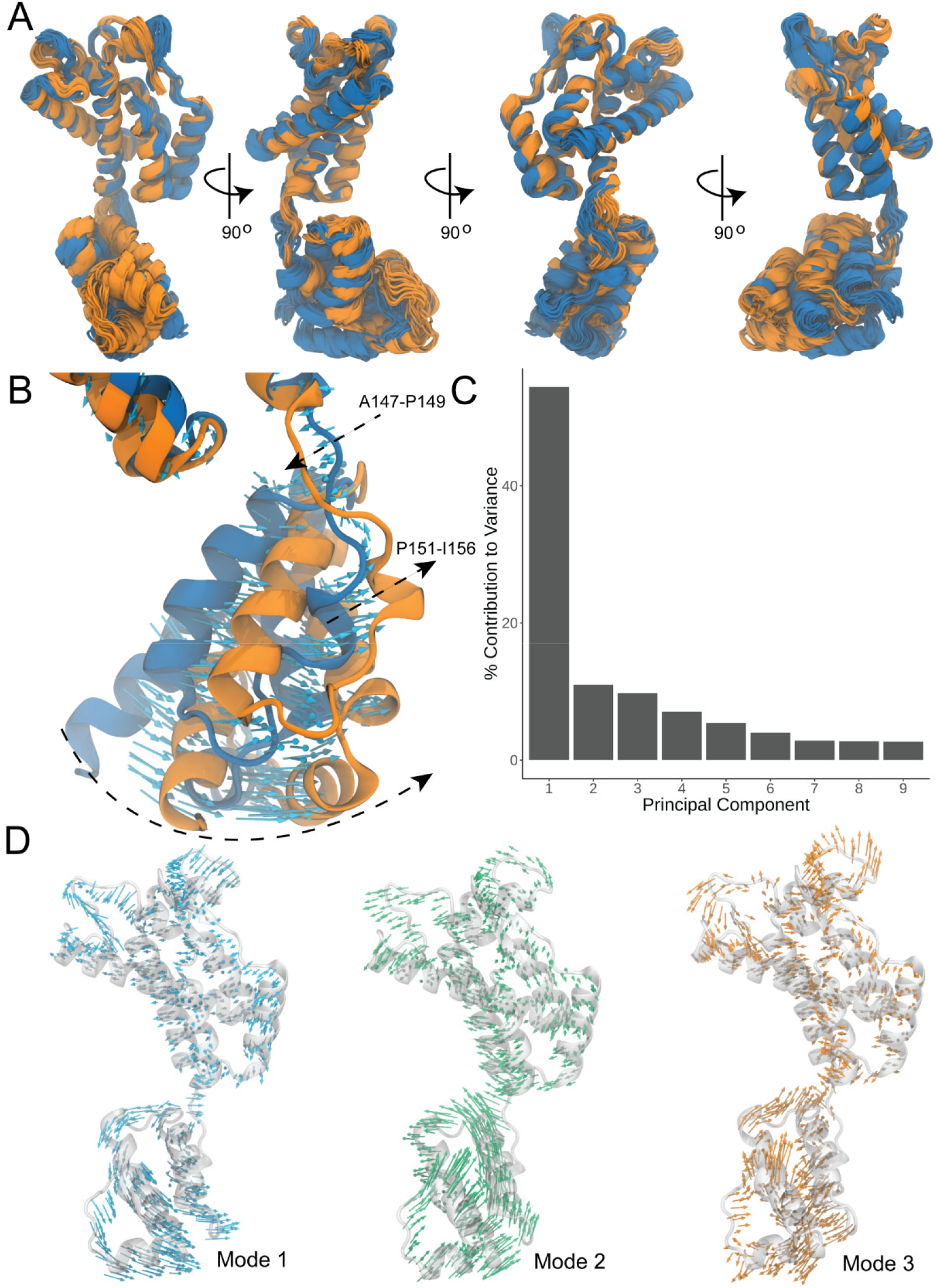
The differences of RSV CA in quasi-equivalent assemblies revealed by PCA. (A). 360° view of CA monomers from the T=1 capsid (orange) aligned by its NTD against that from the tubular (blue) assembly. (B). Principal components reveal a rotation plus translation motion around the flexible interdomain linker to transform a CA in its pentameric assembly to that in the hexameric assembly. The blue vectors denote the direction and magnitude of the motion. (C). Ranking of the contributions of principal components to the differences of the RSV CA conformation in the pentameric vs hexameric assemblies. (D). The residue specific motions of the largest three principal components, shown by colored vectors.

Interestingly, our PCA reveals a dominant mode that shifts an RSV CA monomer in the hexameric tubular assembly to that in the T=1 capsid assembly, contributing to more than half of all the variance in the dataset. As shown in Figures 5B and 5C, this mode corresponds to a 34° rotation and 5.94 Å center-of-mass translation of the CTD mediated by residues in the flexible linker, encompassing residues 147 to 149, as well as residues 151 to 156 in the 3__10__ helix. In fact, nearly the entire structural rearrangements of CTDs can be captured by the top three PCs, as shown in Figure 5D. An animation of the top three PCs is shown in Movie in supporting information. The animation also shows that the additional structural changes in the flexible loop and β-hairpin region are concomitant with the structural rearrangements of the CTD.

The function of the flexible interdomain linker revealed by our work, was hypothesized years ago, by both Pornillos *et al.^16^* and Worthylake *et al.^43^* Brun *et al.^44^* and Wacharapornin *et al.^45^* first demonstrated the detrimental effect of S149A mutation to the HIV viral infectivity. Subsequently, Jiang et al. performed a beautiful systemic site-directed mutagenesis study of the HIV CA flexible linker (residues 145-151). Their results showed that residues in the non-structured linker regions are surprisingly critical to the correct assembly and stability of HIV capsid, with mutations to attenuate or completely abolish infectivity.*^8^* The flexible linker in the HIV CA tubular assemblies adopts four distinct conformations,*^31^* undergoing millisecond scale dynamics by ssNMR,*^46,47^* which would allow such a modulation to take place. Our results support these reports and elucidate the detailed molecular basis to correlate the motions of the flexible linker with the curvature control.

Meanwhile, it is not clear how this motion of flexible linker is triggered and regulated in real viral particle assembly. Solution NMR studies by Lalit *et al.* defined the conformation space of interdomain orientation sampled by HIV CA dimers in the low salt solution, and suggested the selection of the pentameric assembly over hexameric assembly is tightly regulated by such motion.*^3^* Accounting for the conformation space defined by solution NMR, coarse-grained simulations by Xin *et al.* also showed that the HIV CA pentamers can readily be incorporated into hexameric lattice at concentrations comparable to those used in experimental assembly; it demands a highly specific NTD to CTD orientation, not sensitive to particular structural variations within individual domains.*^48^* Subsequent coarse-gained simulations by Grime *et al.* suggest a higher concentration favors the assembly of HIV CA pentamer over hexamer.*^49^* Thus, a possible mechanism for different retroviruses to control their preferred assembly morphology, would be to position accordingly the locations of such factors that promote the specific torsion angles of the interdomain linker suitable for pentameric CA assembly. Slight randomization of their locations would displace the locations of incurred pentamers and give rise to assembly polymorphism.

## Methods

No unexpected or unusually high safety hazards were encountered. Protocols of protein expression and purification, experimental and simulation setup are described in the supporting information.

## Supporting information

Movie showing the PCA analysis to transform a CA monomer from its hexameric state to pentameric state

SupportingInformation4manuscript

## Acknowledgement

The authors acknowledge the CA plasmid from Prof. Rebecca C. Craven at Penn State University and consultation on protein purification and assembly, Woonghee Lee at Univ of Colorado for his help with program NMRSparky, and Frank Shewmaker at Uniform Services University for molecular weight conformation with mass spectrometry. This work was supported by the Mid-career refreshment program of University of Central Florida (B.C.), and the National High Magnetic Field Laboratory through NSF/ DMR-1157490 and the state of Florida. This work was supported by the US National Institutes of Health grants P50AI1504817 (J.R.P.), P20GM104316 (J.R.P.) and R56AI076121. This work used the Extreme Science and Engineering Discovery Environment, which is supported by the National Science Foundation (Grant ACI-1548562) (J.R.P.). This work used XSEDE Bridges and Stampede2 at the Pittsburgh Super Computing Center and Texas Advanced Computing Center, respectively, through allocation MCB170096 (J.R.P.).

## References

[1] Novikova, M., Zhang, Y. L., Freed, E. O., and Peng, K. (2019) Multiple Roles of HIV-1 Capsid during the Virus Replication Cycle, Virologica Sinica 34, 119–134.

[2] Vogt, V. (1997) Retroviral virions and genomes. In Retroviruses, J. M. Coffin, S. H. Hughes, and H.E. Varmus. (New York: Cold Spring Harbor Press), 22–70.

[3] Deshmukh, L., Schwieters, C. D., Grishaev, A., Ghirlando, R., Baber, J. L., and Clore, G. M. (2013) Structure and Dynamics of Full-Length HIV-1 Capsid Protein in Solution, Journal of the American Chemical Society 135, 16133–16147.

[4] Gamble, T. R., Vajdos, F. F., Yoo, S. H., Worthylake, D. K., Houseweart, M., Sundquist, W. I., and Hill, C. P. (1996) Crystal structure of human cyclophilin A bound to the amino-terminal domain of HIV-1 capsid, Cell 87, 1285–1294.

[5] Gitti, R. K., Lee, B. M., Walker, J., Summers, M. F., Yoo, S., and Sundquist, W. I. (1996) Structure of the amino-terminal core domain of the HIV-1 capsid protein, Science 273, 231–235.

[6] Gamble, T. R., Yoo, S. H., Vajdos, F. F., vonSchwedler, U. K., Worthylake, D. K., Wang, H., McCutcheon, J. P., Sundquist, W. I., and Hill, C. P. (1997) Structure of the carboxyl-terminal dimerization domain of the HIV-1 capsid protein, Science 278, 849–853.

[7] Ganser-Pornillos, B. K., von Schwedler, U. K., Stray, K. M., Aiken, C., and Sundquist, W. I. (2004) Assembly properties of the human immunodeficiency virus type 1 CA protein, Journal of Virology 78, 2545–2552.

[8] Jiang, J. Y., Ablan, S. D., Derebail, S., Hercik, K., Soheilian, F., Thomas, J. A., Tang, S. X., Hewlett, I., Nagashima, K., Gorelick, R. J., Freed, E. O., and Levin, J. G. (2011) The interdomain linker region of HIV-1 capsid protein is a critical determinant of proper core assembly and stability, Virology 421, 253–265.

[9] Carnes, S. K., Sheehan, J. H., and Aiken, C. (2018) Inhibitors of the HIV-1 capsid, a target of opportunity, Current Opinion in Hiv and Aids 13, 359–365.

[10] Link, J. O., Rhee, M. S., Tse, W. C., Zheng, J., Somoza, J. R., Rowe, W., Begley, R., Chiu, A., Mulato, A., Hansen, D., Singer, E., Tsai, L. K., Bam, R. A., Chou, C. H., Canales, E., Brizgys, G., Zhang, J. R., Li, J. Y., Graupe, M., Morganelli, P., Liu, Q., Wu, Q. Y., Halcomb, R. L., Saito, R. D., Schroeder, S. D., Lazerwith, S. E., Bondy, S., Jin, D. B., Hung, M., Novikov, N., Liu, X. H., Villasenor, A. G., Cannizzaro, C. E., Hu, E. Y., Anderson, R. L., Appleby, T. C., Lu, B., Mwangi, J., Liclican, A., Niedziela-Majka, A., Papalia, G. A., Wong, M. H., Leavitt, S. A., Xu, Y. L., Koditek, D., Stepan, G. J., Yu, H., Pagratis, N., Clancy, S., Ahmadyar, S., Cai, T. Z., Sellers, S., Wolckenhauer, S. A., Ling, J., Callebaut, C., Margot, N., Ram, R. R., Liu, Y. P., Hyland, R., Sinclair, G. I., Ruane, P. J., Crofoot, G. E., McDonald, C. K., Brainard, D. M., Lad, L., Swaminathan, S., Sundquist, W. I., Sakowicz, R., Chester, A. E., Lee, W. E., Daar, E. S., Yant, S. R., and Cihlar, T. (2020) Clinical targeting of HIV capsid protein with a long-acting small molecule, Nature 584, 614+.

[11] Yeager, M. (2011) Design of in Vitro Symmetric Complexes and Analysis by Hybrid Methods Reveal Mechanisms of HIV Capsid Assembly, Journal of Molecular Biology 410, 534–552.

[12] Ganser, B. K., Li, S., Klishko, V. Y., Finch, J. T., and Sundquist, W. I. (1999) Assembly and analysis of conical models for the HIV-1 core, Science 283, 80–83.

[13] Ganser-Pornillos, B. K., Cheng, A., and Yeager, M. (2007) Structure of full-length HIV-1CA: a model for the mature capsid lattice, Cell 131, 70–79.

[14] Pornillos, O., Ganser-Pornillos, B. K., Kelly, B. N., Hua, Y. Z., Whitby, F. G., Stout, C. D., Sundquist, W. I., Hill, C. P., and Yeager, M. (2009) X-Ray Structures of the Hexameric Building Block of the HIV Capsid, Cell 137, 1282–1292.

[15] Pornillos, O., Ganser-Pornillos, B. K., Banumathi, S., Hua, Y. Z., and Yeager, M. (2010) Disulfide Bond Stabilization of the Hexameric Capsomer of Human Immunodeficiency Virus, Journal of Molecular Biology 401, 985–995.

[16] Pornillos, O., Ganser-Pornillos, B. K., and Yeager, M. (2011) Atomic-level modelling of the HIV capsid, Nature 469, 424–+.

[17] Cardone, G., Purdy, J. G., Cheng, N., Craven, R. C., and Steven, A. C. (2009) Visualization of a missing link in retrovirus capsid assembly, Nature 457, 694–U693.

[18] Hyun, J.-K., Radjainia, M., Kingston, R. L., and Mitra, A. K. (2010) Proton-driven Assembly of the Rous Sarcoma Virus Capsid Protein Results in the Formation of Icosahedral Particles, Journal of Biological Chemistry 285, 15056–15064.

[19] Keller, P. W., Huang, R. K., England, M. R., Waki, K., Cheng, N. Q., Heymann, J. B., Craven, R. C., Freed, E. O., and Steven, A. C. (2013) A Two-Pronged Structural Analysis of Retroviral Maturation Indicates that Core Formation Proceeds by a Disassembly-Reassembly Pathway Rather than a Displacive Transition, Journal of Virology 87, 13655–13664.

[20] Zhao, G., Perilla, J. R., Yufenyuy, E. L., Meng, X., Chen, B., Ning, J., Ahn, J., Gronenborn, A. M., Schulten, K., Aiken, C., and Zhang, P. (2013) Mature HIV-1 capsid structure by cryo-electron microscopy and all-atom molecular dynamics, Nature 497, 643–646.

[21] Mattei, S., Glass, B., Hagen, W. J. H., Krausslich, H. G., and Briggs, J. A. G. (2016) The structure and flexibility of conical HIV-1 capsids determined within intact virions, Science 354, 1434–1437.

[22] Bailey, G. D., Hyun, J.-K., Mitra, A. K., and Kingston, R. L. (2012) A Structural Model for the Generation of Continuous Curvature on the Surface of a Retroviral Capsid, Journal of Molecular Biology 417, 212–223.

[23] Gres, A. T., Kirby, K. A., KewalRamani, V. N., Tanner, J. J., Pornillos, O., and Sarafianos, S. G. (2015) X-ray crystal structures of native HIV-1 capsid protein reveal conformational variability, Science 349, 99–103.

[24] Chen, B., and Tycko, R. (2010) Structural and dynamical characterization of tubular HIV-1 capsid protein assemblies by solid state nuclear magnetic resonance and electron microscopy, Protein Science 19, 716–730.

[25] Han, Y., Ahn, J., Concel, J., Byeon, I.-J. L., Gronenborn, A. M., Yang, J., and Polenova, T. (2010) Solid-State NMR Studies of HIV-1 Capsid Protein Assemblies, Journal of the American Chemical Society 132, 1976–1987.

[26] Byeon, I.-J. L., Hou, G., Han, Y., Suiter, C. L., Ahn, J., Jung, J., Byeon, C.-H., Gronenborn, A. M., and Polenova, T. (2012) Motions on the Millisecond Time Scale and Multiple Conformations of HIV-1 Capsid Protein: Implications for Structural Polymorphism of CA Assemblies, Journal of the American Chemical Society 134, 6455–6466.

[27] Bayro, M. J., Chen, B., Yau, W.-M., and Tycko, R. (2014) Site-Specific Structural Variations Accompanying Tubular Assembly of the HIV-1 Capsid Protein, Journal of Molecular Biology 426, 1109–1127.

[28] Lu, J. X., Bayro, M. J., and Tycko, R. (2016) Major Variations in HIV-1 Capsid Assembly Morphologies Involve Minor Variations in Molecular Structures of Structurally Ordered Protein Segments, Journal of Biological Chemistry 291, 13098–13112.

[29] Zhang, H. L., Hou, G. J., Lu, M. M., Ahn, J., Byeon, I. J. L., Langmead, C. J., Perilla, J. R., Hung, I., Gor’kov, P. L., Gan, Z. H., Brey, W. W., Case, D. A., Schulten, K., Gronenborn, A. M., and Polenova, T. (2016) HIV-1 Capsid Function Is Regulated by Dynamics: Quantitative. Atomic-Resolution Insights by Integrating Magic-Angle-Spinning NMR, QM/MM, and MD, Journal of the American Chemical Society 138, 14066–14075.

[30] Liu, C., Perilla, J. R., Ning, J. Y., Lu, M. M., Hou, G. J., Ramalho, R., Himes, B. A., Zhao, G. P., Bedwell, G. J., Byeon, I. J., Ahn, J., Gronenborn, A. M., Prevelige, P. E., Rousso, I., Aiken, C., Polenova, T., Schulten, K., and Zhang, P. J. (2016) Cyclophilin A stabilizes the HIV-1 capsid through a novel non-canonical binding site, Nature Communications 7.

[31] Lu, M. M., Russell, R. W., Bryer, A. J., Quinn, C. M., Hou, G. J., Zhang, H. L., Schwieters, C. D., Perilla, J. R., Gronenborn, A. M., and Polenova, T. (2020) Atomic-resolution structure of HIV-1 capsid tubes by magic-angle spinning NMR, Nature Structural & Molecular Biology 27, 863–+.

[32] Jeon, J., Qiao, X., Hung, I., Mitra, A. K., Desfosses, A., Huang, D., Gor’kov, P. L., Craven, R. C., Kingston, R. L., Gan, Z. H., Zhu, F. Q., and Chen, B. (2017) Structural Model of the Tubular Assembly of the Rous Sarcoma Virus Capsid Protein, Journal of the American Chemical Society 139, 2006–2013.

[33] Purdy, J. G., Flanagan, J. M., Ropson, I. J., and Craven, R. C. (2009) Retroviral Capsid Assembly: A Role for the CA Dimer in Initiation, Journal of Molecular Biology 389, 438–451.

[34] Bowzard, J. B., Wills, J. W., and Craven, R. C. (2001) Second-site suppressors of Rous sarcoma virus CA mutations: Evidence for interdomain interactions, Journal of Virology 75, 6850–6856.

[35] Takegoshi, K., Nakamura, S., and Terao, T. (2001) C-13-H-1 dipolar-assisted rotational resonance in magic-angle spinning NMR, Chemical Physics Letters 344, 631–637.

[36] Colvin, M. T., Silvers, R., Ni, Q. Z., Can, T. V., Sergeyev, I., Rosay, M., Donovan, K. J., Michael, B., Wall, J., Linse, S., and Griffin, R. G. (2016) Atomic Resolution Structure of Monomorphic A beta(42) Amyloid Fibrils, Journal of the American Chemical Society 138, 9663–9674.

[37] Lu, J. X., Qiang, W., Yau, W. M., Schwieters, C. D., Meredith, S. C., and Tycko, R. (2013) Molecular Structure of beta-Amyloid Fibrils in Alzheimer's Disease Brain Tissue, Cell 154, 1257–1268.

[38] Tuttle, M. D., Comellas, G., Nieuwkoop, A. J., Covell, D. J., Berthold, D. A., Kloepper, K. D., Courtney, J. M., Kim, J. K., Barclay, A. M., Kendall, A., Wan, W., Stubbs, G., Schwieters, C. D., Lee, V. M. Y., George, J. M., and Rienstra, C. M. (2016) Solid-state NMR structure of a pathogenic fibril of full-length human alpha-synuclein, Nature Structural & Molecular Biology 23, 409–415.

[39] Van Melckebeke, H., Wasmer, C., Lange, A., Ab, E., Loquet, A., Bockmann, A., and Meier, B. H. (2010) Atomic-Resolution Three-Dimensional Structure of HET-s(218-289) Amyloid Fibrils by Solid-State NMR Spectroscopy, Journal of the American Chemical Society 132, 13765–13775.

[40] Hardy, E. H., Verel, R., and Meier, B. H. (2001) Fast MAS total through-bond correlation spectroscopy, Journal of Magnetic Resonance 148, 459–464.

[41] Shen, Y., and Bax, A. (2013) Protein backbone and sidechain torsion angles predicted from NMR chemical shifts using artificial neural networks, Journal of Biomolecular Nmr 56, 227–241.

[42] Jorgensen, W. L., Chandrasekhar, J., Madura, J. D., Impey, R. W., and Klein, M. L. (1983) COMPARISON OF SIMPLE POTENTIAL FUNCTIONS FOR SIMULATING LIQUID WATER, Journal of Chemical Physics 79, 926–935.

[43] Worthylake, D. K., Wang, H., Yoo, S. H., Sundquist, W. I., and Hill, C. P. (1999) Structures of the HIV-1 capsid protein dimerization domain at 2.6 angstrom resolution, Acta Crystallographica Section D-Structural Biology 55, 85–92.

[44] Brun, S., Solignat, M., Gay, B., Bernard, E., Chaloin, L., Fenard, D., Devaux, C., Chazal, N., and Briant, L. (2008) VSV-G pseudotyping rescues HIV-1CA mutations that impair core assembly or stability, Retrovirology 5.

[45] Wacharapornin, P., Lauhakirti, D., and Auewarakul, P. (2007) The effect of capsid mutations on HIV-1 uncoating, Virology 358, 48–54.

[46] Byeon, I. J. L., Hou, G. J., Han, Y., Suiter, C. L., Ahn, J., Jung, J., Byeon, C. H., Gronenborn, A. M., and Polenova, T. (2012) Motions on the Millisecond Time Scale and Multiple Conformations of HIV-1 Capsid Protein: Implications for Structural Polymorphism of CA Assemblies, Journal of the American Chemical Society 134, 6455–6466.

[47] Han, Y., Ahn, J., Concel, J., Byeon, I. J. L., Gronenborn, A. M., Yang, J., and Polenova, T. (2010) Solid-State NMR Studies of HIV-1 Capsid Protein Assemblies, Journal of the American Chemical Society 132, 1976–1987.

[48] Qiao, X., Jean, J., Weber, J., Zhu, F. Q., and Chen, B. (2015) Mechanism of polymorphism and curvature of HIV capsid assemblies probed by 3D simulations with a novel coarse grain model, Biochimica Et Biophysica Acta-General Subjects 1850, 2353–2367.

[49] Grime, J. M. A., Dama, J. F., Ganser-Pornillos, B. K., Woodward, C. L., Jensen, G. J., Yeager, M., and Voth, G. A. (2016) Coarse-grained simulation reveals key features of HIV-1 capsid self-assembly, Nature Communications 7.

