## SupportingInformation4manuscript for "Curvature of the retroviral capsid assembly is modulated by a molecular switch"

**Support information for**

### Materials and methods

#### T=1 Spherical RSV CA assembly preparation

The plasmid encoding I190V RSV CA mutant was obtained from Prof. Rebecca C Craven at Penn State University.[1] The plasmid was transformed to *E. coli* strain BL21(DE3) (New England Biolab). The expression and purification of uniform $^{13}C$ and $^{15}N$ labeled I190V CA (U-CA) were completed following a similar protocol as described in our prior work.[2]

Briefly, the transformed bacteria were added into 50 ml 2XYT medium with 50 µl 70 mg/ml Kanamycin and 34 mg/ml Chloramphenicol. The solution was grown at 37°C overnight with 250 rpm shaking. Next morning, 1 Liter minimal medium was inoculated with the overnight culture at initial $OD_{600}$=0.05. Each 1 L minimal medium was prepared with 33.3 ml of 30x salt solution (1 M $Na_2HPO_4$, 1 M $KH_2PO_4$ and 0.1 M $Na_2SO_4$), 2 g of $NH_4Cl$, 4 g of Glucose, 1 ml of 2 M $MgCl_2$, 50 mg of Thiamine, 200 µl of a trace mineral solution (100 mM $FeCl_3$, 1 M $CaCl_2$, 0.5 M $MnCl_2$, 0.5 M $ZnSO_4$, 0.4 M $CoCl_2$, 0.4 M $CuCl_2$, 0.4 M $NiCl_2$, 0.4 M $Na_2MoO_4$, 0.4 M $Na_2SeO_3$ and 0.4 M $H_3BO_3$), and 1 ml of 70 mg/ml Kanamycin and 34 mg/ml Chloramphenicol. With 250 rpm shaking, the 1 Liter culture was then grown to an $OD_{600}$ of 1.0 to 1.5 at 37°C. The cells were spun down by centrifugation at 6000 × g for 10 min and resuspended in a fresh isotopically labeling minimal medium with $^{15}NH_4Cl$ and $^{13}C_6$-D glucose. After 15 min, the expression of U-CA was induced by adding 1 mM isopropyl β-D-1-thiogalactopyranoside (IPTG). The temperature was dropped to 27°C to maximize the yield. After 14 hours, the cells were harvested by centrifuging at 6000 × g.

To lyse the cells, the cell pellets were resuspended in a lysis buffer with 25 mM Tris-HCl at pH 7.5, 50 mM NaCl, 1 mM 4-(2-Aminoethyl) benzenesulfonyl fluoride hydrochloride (AEBSF), 14 µM of Lysozyme and 5 mM β-mercaptoethanol (βME). The solution was incubated on ice with stirring for 1 hr. The dissolved lysis solution was sonicated (Branson Sonifier model 400) at a duty cycle of 20% and an amplitude of for 15 minutes while stirring on ice. This operation was repeated 7 times. After sonication was completed, the solution was centrifuged at 20,000 × g for 1 hr. The cell debris was discarded. The protein in the supernatant was precipitated by addition of 35 %(v/v) of a saturated $(NH_4)_2SO_4$ solution. The solution turned murky and was incubated on ice for 1 hr. The protein pellet was spun down by centrifugation at 20,000 × g for 1 hr. The pellet was then resuspended in 100 ml Sepharose A buffer (25 mM MOPS-KOH with βME at pH 6.9). The solution was desalted by dialysis against 6 L of Sepharose A buffer. It was then subject to 14,000 × g for 15 min and filtered before loading into a Sepharose fast flow (SP FF) column. The protein was eluted by running a linear gradient of Sepharose B buffer (25 mM MOPS-KOH with 1 M NaCl and βME at pH 6.9) over 200 ml. Subsequently the eluted protein was dialyzed against the S75 buffer (10 mM Tris, 50 mM NaCl, pH 8.0 and 5 mM βME). The dialyzed protein solution was then concentrated down to 1.5 ml for running the size-exclusion column (Superdex 75). The typical yield is about 2 mol per Liter. The identity of the final product was confirmed by mass spectrometry.

To promote T=1 spherical assembly, I190V mutant RSV CA was concentrated to 6 mg/ml (estimated from its UV absorption at 280 nm) and then mixed with equal volume of 1 M $Na_2HPO_4$ / $NaH_2PO_4$ buffer pH 8.0. The tube containing the solution was flipped a few times to homogenize the mixture. The solution was then left undisturbed for about 3 days for the assembly to complete. Transmission electron microscopy was applied to evaluate the identity of the assembly and the assembled material was pelleted by centrifugation at 14,000 × g for 1 hr. The pelleted assembly was packed into 3.2 mm rotors for solid-state NMR measurements. The integrity of the assembly was evaluated by comparing 1D $^{13}C$ CP NMR spectra before and after long sessions of 3D NCaCx/NCOCx experiments.

**Transmission electron microscopy**

The I190V RSV CA assembly morphologies were verified with a JEOL 1011 transmission electron microscope operated at 80 keV. To prepare a sample, 5 μL of the assembly solution was pippetted onto a holey carbon TEM grid (part no. 01824, Ted Pella inc.). The solution was left on the grid for 5 min before removing excess solution by blotting. 2 μL of 3 %(w/v) uranyl acetate was applied to the grid, which was immediately blotted, rinsed with 2 μL of filtered DI water, and finally blotted dry for TEM imaging.

**Solid-state NMR**

Multidimensional NMR spectra were acquired on an Agilent spectrometer operating at $^{1}H$ NMR frequency of 600 MHz (14.1 Tesla) equipped with a 3.2 mm BioMAS probe at University of Central Florida and on Bruker spectrometers with $^{1}H$ NMR frequency of 600 and 800 MHz equipped with 3.2 mm E-free probes in the National High Magnetic Field Laboratory (NHMFL). All spectra for resonance assignments were acquired at 13.5 kHz Magic Angle Spinning (MAS) frequency with SPECIFIC-CP (cross polarization) for $^{15}N$-$^{13}C$ transfer and DARR (dipolar assisted rotational resonance) for homonuclear $^{13}C$-$^{13}C$ mixing. All spectra were processed with NMRpipe.[3] Each dimension was zero-filled to twice the acquisition number. Then a Gaussian broadening function was applied to the direct and indirect dimension. Forward linear prediction was applied with twice the number of data points in indirect dimensions. Details of the processing parameters are detailed in Table S1 in the supporting information. The spectra were plotted in Sparky,[4] with the floor contour level shown in Table S1.

**Simulation overview**

The data-guided RSV T=1 capsids were simulated at a constant temperature of 310 K and pressure of 1 atm, with 2-fs time steps, using the NAMD molecular dynamics engine[5] and the CHARMM36m protein force field.[6] Dihedral angle restraints, derived from experimental NMR data, were generated with the TALOS-N software[7] and incorporated into the extraBonds module in NAMD. Molecular dynamics flexible fitting (MDFF) was accomplished with the gridForces module[8] in NAMD. Each complete RSV T=1 capsid system comprised 12 CA pentamers, or 60 RSV monomers, in total and contained 1,821,665 atoms including solvent and ions.

**System preparation**

Prior to constructing the RSV T=1 capsid, protonation states of RSV CA were assigned using propKa software[9] at pH 7.0. A short energy minimization – 500 steps – of the RSV CA monomer with NMR restraints was performed with NAMD. The CA monomer was then rigid-body docked into the 8.5 Å cryogenic electron microscopy (cryo-EM) density[10] using Chimera[11]. Next, the T=1 assembly was constructed by applying a transformation matrix derived from a T=1 icosahedral symmetry to the coordinates of the monomer. Afterwards, the net charge of the T=1 assembly was calculated as +120e, and 120 Chloride counter ions were placed according to iterative calculations of the Coulombic potential, using the cionize tool in VMD,[12] with a specified grid spacing of 1.0 Å.

Solvation of the protein and counter ions was performed using the Solvate tool in VMD. The dimensions of water box were determined to be 230x230x230 Å, which allowed at least 5.0 Å of padding from the protein along each Cartesian axis while exceeding the bounds of the EM density maps, the latter a requirement for MDFF.[8] Addition of bulk Na+ and Cl- ions to a final salt concentration of 500 mM was carried out using the AutoIonize VMD plugin. After solvation and addition of bulk ions, each system totaled 1.8 million atoms. The latter were parametrized using the CHARMM36m protein parameter set.

Prior to production NMR-guided MDFF simulations, each system was subjected to 50,000 cycles of steepest descent minimization in NAMD with protein atoms restrained to allow the water box to minimize. Then, 20,000 cycles of minimization with only backbone atom restraints and 20,000 of unrestrained minimization were performed. For the latter 20,000 cycles of minimization, the gridForce module in NAMD was employed with a force constant of 0.10 kcal*mol$^{-1}$*Å$^{-2}$, with each replica using its respective EM density.

### Integration of experimental data

NMR dihedral angles were introduced as extraBonds within NAMD. The force constant for dihedral angles was 0.6 kcal*mol$^{-1}$*Å$^{-2}$. Grid forces to perform MDFF were applied with the gridForces module in NAMD,[5] using a force constant of 0.15 kcal*mol$^{-1}$*Å$^{-2}$ during production.

### Molecular dynamics simulations

Molecular dynamics simulations were performed on the Los Alamos National Laboratory's (LANL) Grizzly supercomputer, using the NAMD molecular dynamics engine[5] and the CHARMM36m protein force field.[6] For non-bonded interactions, atom pairs within one, two or three bonds of each other were excluded. A cutoff of 12.0 Å was employed, using a switching distance of 10.0 Å to smooth the interaction potential beyond the cutoff distance. The maximum distance per pair was 14.0 Å. Full electrostatics were considered using the particle mesh Ewald (PME) method[13, 14] with a grid spacing of 2.0 Å and eighth-order interpolation. Short range non-bonded interactions were evaluated every 2.0 fs and full electrostatics every 4.0 fs. Electrostatic forces were split between short and long range potentials using a quintic polynomial splitting function.[15] Pressure was maintained with the Nose–Hoover Langevin barostat, using a target pressure of 1.01325 bar and target temperature of 310 K; the specified oscillation period was 20 ps with a decay period of 10 ps. Temperature was maintained at 310 K using stochastic velocity rescaling[16] and a rescale period of 200 fs. A time step of 2.0 fs per step was used throughout the simulation and all bonds to hydrogen were constrained with the RATTLE and SHAKE algorithms for solvent and solute, respectively.[17, 18]

Table S1. Spectra acquired in this work with Uniform $^{13}C$, $^{15}N$ labeled I190V RSV CA.

| Spectrum | Field (MHz) in 1H frequency | Recoupling | Process Parameter | | | Contour noise level |
|---|---|---|---|---|---|---|
| | | | Linear prediction | Apodization | Zero-filling | |
| 2D $^{13}C$-$^{13}C$ correlation | 600 | 20 ms DARR | 212 | G1 60, g2 90 | Auto | 3e+6 |
| 2D NCaCx | 800 | 50 ms DARR | 212 | G1 90, G2 90 | Auto | 1e+7 |
| 2D NCOCx | 800 | 150 ms DARR | 212 | G1 90, G2 90 | Auto | 6e+6 |
| 3D NCaCx | 800 | 50 ms DARR | 92 | G1 0, G2 60, C 1.0 | 4096 | 4e+7 |
| 3D NCOCx | 800 | 150 ms DARR | 98 | G1 0, G2 30, G3 0.1, C 1.0 | 4096 | 4.4e+8 |

Table S2. Sequential assignments of the T=1 capsid assembly of the I190V RSV CA mutant. Resonances corresponding to residues with a large difference in torsion angle between T=1 capsid and tubular assemblies are highlighted in green. Residues assigned from the TOBSY experiment highlighted in blue. Residues not assigned are highlighted in red.

| Res. Num. | Res. Type | N | Ca | CO | Cb | Cg1/Cg | Cg2 | Cd1/Cd | Cd2 | Ce/Cz | Aromatic |
|---|---|---|---|---|---|---|---|---|---|---|---|
| 1 | P | - | 61.8 | 171.4 | 32.09 | 26.17 | | 48.58 | | | |
| 2 | V | 121.9 | 61.7 | 175.2 | 34.69 | 22.45 | 21.69 | | | | |
| 3 | V | 125.5 | 59.3 | 173.7 | 34.81 | 20.75 | 20.26 | | | | |
| 4 | I | 125.5 | 60.8 | 176 | 35.61 | 26.93 | 17.76 | 10.71 | | | |
| 5 | K | 128.8 | 54.6 | 177.5 | 33.93 | 25.36 | | 28.69 | | - | |
| 6 | T | 117.7 | 58.8 | 177.916 | 65.53 | 24.312 | | | | | |
| 7 | E | 125.8 | 54.6 | 176.5 | 31.94 | 35.87 | | 184.1 | | | |
| 8 | G | 106.6 | 44.2 | 170.974 | | | | | | | |
| 9 | P | 135.6 | 63.2 | 171.7 | 32.47 | 27.91 | | 49.71 | | | |
| 10 | A | 122.3 | 52.3 | 176.9 | 19.47 | | | | | | |
| 11 | W | 121.5 | 56.6 | 176 | 28.27 | | | | | | 109.1 |
| 12 | T | 124.9 | 57.1 | 170 | 70.8 | 20.4 | | | | | |
| 13 | P | 139.5 | 62.9 | 177.159 | 33.93 | 26.614 | | 51.882 | | | |
| 14 | L | 122.5 | 54.8 | 176.494 | 42.76 | 27.35 | | 25.93 | 23.44 | | |
| 15 | E | 123.7 | 54.3 | 176.311 | 29.73 | 36.37 | | 180.3 | | | |
| 16 | P | 139.5 | 64.4 | 177.587 | 32.287 | 27.43 | | 51.062 | | | |
| 17 | K | 116.4 | 59.5 | 179.3 | 33.09 | 26.46 | | 32.033 | | - | |
| 18 | L | 119.1 | 58.2 | 178.435 | 41.968 | 28.24 | | 25.814 | 22.641 | | |
| 19 | I | 120.9 | 59.6 | 178 | 37.56 | 29.51 | 18.93 | 9.047 | | | |
| 20 | T | 117.2 | 67.1 | 176 | 67.85 | 20.91 | | | | | |
| 21 | R | 121.3 | 59.4 | 179 | 29.97 | 27.6 | | 43.42 | | 151.5 | |
| 22 | L | 121.9 | 57.8 | 178.095 | 41.75 | 27.259 | | 26.29 | 23.237 | | |
| 23 | A | 120.8 | 55.3 | 180.237 | 19.04 | | | | | | |
| 24 | D | 119.7 | 57.2 | 178.6 | 40.41 | 181 | | | | | |
| 25 | T | 122.9 | 66.9 | 175.9 | 67.96 | 17.89 | | | | | |
| 26 | V | 121.9 | 66.8 | 179.5 | 31.59 | 22.65 | 21.05 | | | | |
| 27 | R | 118.5 | 59.0 | 176.9 | 30.42 | 27.77 | | 35.6 | | 154.1 | |
| 28 | T | 107.8 | 63.6 | 176.5 | 70.11 | 21.768 | | | | | |
| 29 | K | 119.9 | 55.2 | 178 | 32.62 | 25.08 | | 28.05 | | 42.43 | |
| 30 | G | 108.6 | 44.4 | 173.4 | | | | | | | |
| 31 | L | 120.4 | 55.4 | 177.6 | 42.43 | 28.32 | | 26.7 | 23.73 | | |
| 32 | R | 113.1 | 55.6 | 176.8 | 31.48 | 29.05 | | 44.35 | | - | |
| 33 | S | 114.1 | 55.0 | 173.859 | 64.171 | | | | | | |
| 34 | P | 132.5 | 65.8 | 177.3 | 30.81 | 27.39 | | 49.77 | | | |
| 35 | I | 132.9 | 62.6 | 176.4 | 38.21 | 27.99 | 17.55 | 11.68 | | | |
| 36 | T | 118.8 | 66.6 | 176.1 | 68.44 | 22.04 | | | | | |
| 37 | M | 118.0 | 57.2 | 178.2 | 32.04 | 31.06 | | | | 17.238 | |
| 38 | A | 120.3 | 55.2 | 180.2 | 17.96 | | | | | | |
| 39 | E | 119.6 | 60.1 | 179.462 | 28.54 | 34.131 | | 181.4 | | | |
| 40 | V | 120.5 | 67.1 | 178.1 | 31.95 | 22.67 | 20.83 | | | | |
| 41 | E | 119.7 | 60.1 | 179.462 | 28.54 | 34.131 | | 182.5 | | | |
| 42 | A | 119.7 | 54.9 | 178.7 | 17.96 | | | | | | |
| 43 | L | 120.4 | 57.8 | 176.997 | 42.037 | 27.178 | | 24.653 | 21.74 | | |
| 44 | M | 113.9 | 55.3 | 174.8 | 31.98 | 31.22 | | | | 18.19 | |
| 45 | S | 113.1 | 60.3 | 174.7 | 63.35 | | | | | | |
| 46 | S | 118.7 | 56.0 | 174.9 | 63.85 | | | | | | |
| 47 | P | - | - | - | - | - | | - | | | |

| | | | | | | | | | | | |
|---|---|---|---|---|---|---|---|---|---|---|---|
| 48 | L | 124.3 | 53.3 | 174.057 | 44.399 | 26.216 | | 24.28 | 23.304 | | |
| 49 | L | 111.8 | 52.5 | 177.4 | 41.14 | 28.41 | | 26.26 | 23.35 | | |
| 50 | P | - | - | - | - | - | | - | | | |
| 51 | H | 116.6 | 60.3 | 178 | 31.31 | | | | | | - |
| 52 | D | 116.5 | 56.8 | 174 | 38.86 | 179.4 | | | | | |
| 53 | V | 115.1 | 66.1 | 176.9 | 31.72 | 23.43 | 22.41 | | | | |
| 54 | T | 112.3 | 65.7 | 176.947 | 67.7 | 23.03 | | | | | |
| 55 | N | 119.8 | 56.6 | 175.2 | 40.36 | 176.743 | | | | | |
| 56 | L | 120.7 | 57.8 | 176.995 | 39.275 | 26.944 | | 25.93 | 24.65 | | |
| 57 | M | 114.7 | 56.4 | 178.4 | 32.07 | 30.35 | | | | 17.71 | |
| 58 | R | 118.5 | 59.0 | 176.9 | 30.43 | 27.77 | | - | | - | |
| 59 | V | 115.3 | 65.5 | 178.205 | 31.843 | 22.776 | 21.789 | | | | |
| 60 | I | 109.4 | 63.3 | 175.7 | 38.74 | 25.96 | 17.64 | 13.75 | | | |
| 61 | L | 119.4 | 53.3 | 177.5 | 43.75 | 27.58 | | 26.27 | 23.26 | | |
| 62 | G | 108.9 | 44.1 | 171.4 | | | | | | | |
| 63 | P | 135.0 | 64.6 | 177.8 | 32.6 | 27.1 | | 53.32 | | | |
| 64 | A | 119.2 | 56.8 | 176.2 | 15.21 | | | | | | |
| 65 | P | 132.4 | 65.8 | 176.3 | 30.81 | 27.39 | | 49.77 | | | |
| 66 | Y | 118.6 | 61.7 | 177.8 | 39.38 | | | | | | 130.3 |
| 67 | A | 122.7 | 55.2 | 177.034 | 18.07 | | | | | | |
| 68 | L | 118.2 | 56.6 | 178.9 | 43.05 | 26.88 | | 25.16 | - | | |
| 69 | W | 121.5 | 56.6 | 175.964 | 28.27 | | | | | | 109.1 |
| 70 | M | 119.9 | 57.5 | 179.6 | 32 | 30.83 | | | | 27.18 | |
| 71 | D | 119.7 | 56.7 | 176.745 | 40.36 | 179.704 | | | | | |
| 72 | A | 122.3 | 55.3 | 178 | 18.07 | | | | | | |
| 73 | W | 123.3 | 59.6 | 178.2 | 26.35 | | | | | | 122.1 |
| 74 | G | 104.8 | 48.1 | 175.6 | | | | | | | |
| 75 | V | 121.9 | 66.8 | 179.5 | 31.59 | 22.65 | 21.05 | | | | |
| 76 | Q | 119.6 | 60.1 | 177.4 | 28.54 | 34.13 | | 179.5 | | | |
| 77 | L | 120.8 | 57.8 | 179.828 | 39.278 | 26.944 | | 25.926 | 21.739 | | |
| 78 | Q | 117.9 | 60.2 | 176 | 25.12 | 32.44 | | 183.1 | | | |
| 79 | T | 117.8 | 66.1 | 176.6 | 67.97 | 17.91 | | | | | |
| 80 | V | 125.3 | 65.9 | 177.2 | 30.65 | 21.95 | 20.43 | | | | |
| 81 | I | 118.9 | 62.9 | 179.193 | 35.48 | 27.994 | 17.573 | 11.42 | | | |
| 82 | A | 124.5 | 55.5 | 180.3 | 17.47 | | | | | | |
| 83 | A | 122.4 | 55.3 | 179.6 | 18.07 | | | | | | |
| 84 | A | 121.5 | 53.4 | 178.1 | 17.13 | | | | | | |
| 85 | T | 114.5 | 65.2 | 174.9 | 68.75 | 20.69 | | | | | |
| 86 | R | 120.0 | 57.7 | 175.8 | 30.82 | 27.18 | | 43.11 | | - | |
| 87 | D | 116.1 | 49.7 | 174 | 42.23 | 180.4 | | | | | |
| 88 | P | - | - | - | - | - | | - | | | |
| 89 | R | 115.2 | 55.5 | 175.3 | 29.09 | 27.27 | | 43.03 | | 144.2 | |
| 90 | H | 124.7 | 56.9 | 169.984 | 31.97 | | | | | | 138 |
| 91 | P | 131.6 | 65.5 | 181.898 | 30.643 | 24.389 | | 49.41 | | | |
| 92 | A | 122.2 | 55.2 | 178.04 | 18.85 | | | | | | |
| 93 | N | 117.8 | 54.2 | 176.5 | 40.93 | 177.5 | | | | | |
| 94 | G | 108.6 | 44.4 | 173.4 | | | | | | | |
| 95 | Q | - | - | - | - | - | | - | | | |
| 96 | G | 108.8 | 44.0 | 176.048 | | | | | | | |
| 97 | R | 120.1 | 57.8 | 177 | 30.82 | 27.18 | | 43.12 | | - | |
| 98 | G | 112.9 | 49.6 | 171.8 | | | | | | | |
| 99 | E | 119.6 | 56.8 | 176.746 | 30.86 | 40.36 | | 181.1 | | | |
| 100 | R | 120.4 | 55.4 | 177.4 | 28.32 | 26.7 | | 42.43 | | - | |
| 101 | T | 118.0 | 63.1 | 172.6 | 65.94 | 19.1 | | | | | |
| 102 | N | 113.7 | 53.1 | 174.1 | 39.88 | 175.8 | | | | | |

| | | | | | | | | | | | |
|---|---|---|---|---|---|---|---|---|---|---|---|
| 103 | L | 120.4 | 58.1 | 176.996 | 42.03 | 26.38 | | 25.21 | 24.14 | | |
| 104 | N | 115.3 | 55.8 | 178.3 | 36.6 | 175.3 | | | | | |
| 105 | R | 120.6 | 60.6 | 177.695 | 30.77 | 29.28 | | 43.87 | | 159.9 | |
| 106 | L | 114.1 | 56.9 | 175 | 41.22 | 26.92 | | 25.88 | 21.634 | | |
| 107 | K | 108.1 | 56.1 | 177.816 | 34.177 | 26.086 | | 29.644 | | 46.864 | |
| 108 | G | 108.6 | 46.8 | 176.478 | | | | | | | |
| 109 | L | 116.0 | 53.4 | 176.6 | 43.153 | 25.5 | | 22.208 | 19.49 | | |
| 110 | A | 121.4 | 51.3 | 175.2 | 19.41 | | | | | | |
| 111 | D | 119.7 | 56.7 | 176.746 | 40.36 | 179.703 | | | | | |
| 112 | G | 112.5 | 43.8 | 175.1 | | | | | | | |
| 113 | M | 113.5 | 56.8 | 177.325 | 34.19 | 33.8 | | | | 21.63 | |
| 114 | V | 121.4 | 65.0 | 177.2 | 30.53 | 22.29 | 20.69 | | | | |
| 115 | G | 118.8 | 46.8 | 174.2 | | | | | | | |
| 116 | N | 112.9 | 49.5 | 171.8 | 38.16 | 179 | | | | | |
| 117 | P | - | - | - | - | - | | - | | | |
| 118 | Q | 116.4 | 59.5 | 177.7 | 26.457 | 32.03 | | 179.3 | | | |
| 119 | G | 108.8 | 47.1 | 175.1 | | | | | | | |
| 120 | Q | 119.7 | 60.1 | 177.406 | 28.541 | 34.128 | | 179.492 | | | |
| 121 | A | 118.8 | 54.4 | 177.7 | 18.29 | | | | | | |
| 122 | A | 117.5 | 53.7 | 177.8 | 18.95 | | | | | | |
| 123 | L | 115.7 | 56.2 | 179.282 | 45.61 | 27.07 | | 25.183 | 22.36 | | |
| 124 | L | 120.4 | 55.4 | 177.268 | 42.432 | 28.315 | | 26.709 | 23.728 | | |
| 125 | R | 123.5 | 54.3 | 176.311 | 29.94 | 27.39 | | 42.95 | | 159.1 | |
| 126 | P | 137.7 | 66.6 | 175 | 32.04 | 27.41 | | 50.51 | | | |
| 127 | G | 102.8 | 46.4 | 178.6 | | | | | | | |
| 128 | E | 119.8 | 60.1 | 177.4 | 28.54 | 33.29 | | 183.1 | | | |
| 129 | L | 117.4 | 58.5 | 181.05 | 41.024 | 26.602 | | 25.744 | 24.313 | | |
| 130 | V | 120.5 | 67.1 | 178.098 | 31.954 | 22.672 | 20.831 | | | | |
| 131 | A | 122.3 | 55.2 | 181.3 | 18.85 | | | | | | |
| 132 | I | 123.1 | 66.8 | 178.9 | 39.54 | 30.62 | 17.98 | - | | | |
| 133 | T | 116.4 | 66.2 | 177.3 | 68.32 | 21.71 | | | | | |
| 134 | A | 124.6 | 55.7 | 182 | - | | | | | | |
| 135 | S | 113.9 | 62.6 | 176 | 63.96 | | | | | | |
| 136 | A | 125.7 | 55.8 | 179.6 | 20.39 | | | | | | |
| 137 | L | 121.9 | 57.8 | 178.094 | 41.75 | 27.258 | | 26.291 | 23.236 | | |
| 138 | Q | 119.8 | 55.2 | 175.9 | 28.05 | 32.33 | | 182.1 | | | |
| 139 | A | 125.7 | 55.8 | 179.6 | 20.39 | | | | | | |
| 140 | F | 119.4 | 63.2 | 175.9 | 38.78 | | | | | | 131.1 |
| 141 | R | 118.0 | 60.3 | 176.7 | 32.44 | 25.122 | | 43.559 | | 180.3 | |
| 142 | E | 118.7 | 59.2 | 179.3 | 27.77 | 31.22 | | 182.9 | | | |
| 143 | V | 121.8 | 66.8 | 179.5 | 31.6 | 22.65 | 21.05 | | | | |
| 144 | A | 122.1 | 55.3 | 178 | 16.73 | | | | | | |
| 145 | R | 113.4 | 58.6 | 177.8 | 30.13 | 27.97 | | 43.06 | | - | |
| 146 | L | 126.8 | 62.3 | 171.7 | 48.85 | 32.09 | | 26.37 | 22.54 | | |
| 147 | A | 122.3 | 52.3 | 176.9 | 19.47 | | | | | | |
| 148 | E | 136.1 | 49.9 | 175.6 | 27.81 | 32.13 | | 179.2 | | | |
| 149 | P | 135.5 | 64.2 | 176.9 | 32.6 | 27.1 | | 50.74 | | | |
| 150 | A | 123.4 | 53.4 | 175.1 | 18.02 | | | | | | |
| 151 | G | 113.0 | 44.5 | 174.5 | | | | | | | |
| 152 | P | 135.7 | 63.1 | 176.3 | 32.47 | 27.91 | | 49.71 | | | |
| 153 | W | 119.7 | 60.1 | 177.4 | 27.5 | | | | | | 114.3 |
| 154 | A | 122.3 | 55.2 | 181.3 | 18.07 | | | | | | |
| 155 | D | 117.7 | 54.2 | 176.523 | 40.93 | 181.2 | | | | | |
| 156 | I | 121.5 | 66.3 | 177.5 | 37.76 | 30.48 | 18.83 | 14.72 | | | |
| 157 | M | - | - | - | - | - | | | | - | |

| | | | | | | | | | | | |
|---|---|---|---|---|---|---|---|---|---|---|---|
| 158 | Q | 126.3 | 56.1 | 177.2 | 27.18 | 32.05 | | 178.3 | | | |
| 159 | G | 119.1 | 45.0 | 172.9 | | | | | | | |
| 160 | P | 134.7 | 64.8 | 177.8 | 31.37 | 27.56 | | 49.75 | | | |
| 161 | S | 113.0 | 56.7 | 172.794 | 63.49 | | | | | | |
| 162 | E | 125.8 | 54.6 | 176.5 | 31.94 | 35.87 | | 184.1 | | | |
| 163 | S | 127.7 | 58.1 | 174.7 | 63.92 | | | | | | |
| 164 | F | 124.3 | 63.1 | 177.1 | 38.88 | | | | | | - |
| 165 | V | 117.2 | 67.0 | 176 | 31.55 | 23.76 | 20.91 | | | | |
| 166 | D | 120.9 | 57.9 | 179.8 | 39.28 | 178.2 | | | | | |
| 167 | F | 124.0 | 59.1 | 175.8 | 38.69 | | | | | | 130.5 |
| 168 | A | 122.3 | 55.0 | 177.216 | 18.07 | | | | | | |
| 169 | N | 116.1 | 56.4 | 179.3 | 37.46 | 182.6 | | | | | |
| 170 | R | 120.2 | 60.4 | 177.695 | - | - | | - | | - | |
| 171 | L | 121.9 | 57.8 | 178.1 | 41.75 | 27.261 | | 26.289 | 23.238 | | |
| 172 | I | 119.2 | 65.5 | 176.5 | 38.35 | 31.13 | - | - | | | |
| 173 | K | - | - | - | - | - | | - | | - | |
| 174 | A | 120.0 | 55.1 | 177.6 | 17.96 | | | | | | |
| 175 | V | 117.2 | 67.1 | 176 | 31.55 | 24.25 | 20.91 | | | | |
| 176 | E | 120.5 | 59.9 | 177.2 | 28.54 | 37.56 | | 182.7 | | | |
| 177 | G | 103.7 | 45.2 | 173.6 | | | | | | | |
| 178 | S | 117.9 | 59.1 | 173.546 | 65.53 | | | | | | |
| 179 | D | 117.7 | 54.3 | 176.5 | 40.78 | 181.2 | | | | | |
| 180 | L | 122.4 | 54.8 | 176.5 | 42.76 | 27.35 | | 25.93 | 23.44 | | |
| 181 | P | 136.4 | 66.6 | 178.223 | 32.081 | 27.949 | | 50.017 | | | |
| 182 | P | 135.9 | 66.2 | 179.2 | 32.08 | 27.74 | | 50.1 | | | |
| 183 | S | 110.9 | 60.2 | 174.7 | 62.33 | | | | | | |
| 184 | A | 121.4 | 51.3 | 179.6 | 19.41 | | | | | | |
| 185 | R | 115.5 | 61.0 | 176.3 | 30.2 | 27.46 | | 44.25 | | 160.8 | |
| 186 | A | 119.2 | 56.8 | 176.2 | 15.21 | | | | | | |
| 187 | P | 131.8 | 65.6 | 181.9 | 31.04 | - | | 49.41 | | | |
| 188 | V | 117.3 | 67.0 | 179.7 | 31.55 | 23.76 | 20.91 | | | | |
| 189 | I | 119.8 | 57.4 | 180.4 | 40.41 | 30.83 | 27.108 | 17.48 | | | |
| 190 | I | 119.5 | 70.0 | 176.4 | 38.62 | 24.74 | 21.71 | | | | |
| 191 | D | 119.7 | 57.5 | 175.774 | 40.41 | 179.604 | | | | | |
| 192 | C | 118.2 | 56.6 | 177.5 | 26.87 | | | | | | |
| 193 | F | 119.4 | 63.2 | 175.9 | 38.78 | | | | | | 131.1 |
| 194 | R | 116.3 | 60.1 | 178.449 | 31.08 | 28.793 | | 43.49 | | - | |
| 195 | Q | 112.4 | 56.7 | 176.8 | 31.29 | 33.42 | | 178.2 | | | |
| 196 | K | 115.2 | 55.6 | 177.6 | 36.6 | 27.27 | | 29.07 | | 43.03 | |
| 197 | S | 113.1 | 56.7 | 172.794 | 64.81 | | | | | | |
| 198 | Q | 119.8 | 55.0 | 175.9 | 28.05 | 32.62 | | 178.3 | | | |
| 199 | P | 136.7 | 66.8 | 178.2 | 31.26 | 27.79 | | 50.02 | | | |
| 200 | D | 116.4 | 56.7 | 178.929 | 38.86 | 173.98 | | | | | |
| 201 | I | 123.8 | 62.3 | 178.3 | 35.61 | 28.28 | 18.14 | 10.11 | | | |
| 202 | Q | 119.7 | 60.1 | 177.4 | 28.54 | 34.51 | | 181.4 | | | |
| 203 | Q | 115.8 | 60.9 | 179.2 | 27.46 | 30.43 | | 183.8 | | | |
| 204 | L | 120.4 | 58.1 | 176.996 | 42.034 | 26.38 | | 25.207 | 24.139 | | |
| 205 | I | 119.0 | 65.8 | 176.2 | 37.72 | 30.66 | 19.84 | 15.52 | | | |
| 206 | R | 119.4 | 60.2 | 179.463 | 31.854 | 27.38 | | 42.38 | | 161.7 | |
| 207 | T | 125.3 | 65.9 | 177.2 | 68.33 | 21.95 | | | | | |
| 208 | A | 125.6 | 50.6 | 175.7 | 22.22 | | | | | | |
| 209 | P | 135.7 | 63.1 | 176.3 | 32.47 | 27.91 | | 49.71 | | | |
| 210 | S | 117.9 | 59.1 | 173.538 | 65.53 | | | | | | |
| 211 | T | 117.2 | 68.1 | 175.3 | 68.86 | 22.44 | | | | | |
| 212 | L | 124.3 | 53.4 | 174.1 | 44.4 | - | | 26.22 | 24.28 | | |

| | | | | | | | | | | | |
|---|---|---|---|---|---|---|---|---|---|---|---|
| 213 | T | 112.3 | 62.6 | 174.3 | 70.72 | 21.39 | | | | | |
| 214 | T | 111.0 | 58.2 | 173.2 | 68.85 | 21.77 | | | | | |
| 215 | P | 132.5 | 65.7 | 179.5 | 31.08 | 27.39 | | 49.77 | | | |
| 216 | G | 103.1 | 46.5 | 178.4 | | | | | | | |
| 217 | E | 119.6 | 60.1 | 179.462 | 28.54 | 34.131 | | 181.4 | | | |
| 218 | I | - | - | - | - | - | - | - | | | |
| 219 | I | - | - | - | - | - | - | - | | | |
| 220 | K | 117.5 | 59.8 | 179.8 | 32.82 | 24.97 | | 29.53 | | 42.13 | |
| 221 | Y | 118.4 | 62.1 | 177.3 | 38.94 | | | | | | 130.6 |
| 222 | V | 118.2 | 66.0 | 176.8 | 30.34 | 18.96 | 17.9 | | | | |
| 223 | L | 118.9 | 58.2 | 178.1 | 41.97 | 26.827 | | 25.813 | 22.641 | | |
| 224 | D | 119.7 | 56.7 | 176.746 | 40.36 | 179.704 | | | | | |
| 225 | R | 118.2 | 56.6 | 178.7 | 26.88 | 25.16 | | 43.29 | | - | |
| 226 | Q | 118.7 | 59.1 | 179.3 | 29.15 | 35.6 | | 182.9 | | | |
| 227 | K | 119.9 | 57.9 | 175.8 | 30.83 | 24.14 | | 28.23 | | 43.11 | |
| 228 | T | 114.2 | 55.0 | 173.859 | 64.17 | - | | | | | |
| 229 | A | - | - | - | - | | | | | | |
| 230 | P | - | - | - | 31.27 | 27.7 | | - | | | |
| 231 | L | - | 57.4 | 173 | 43.78 | 27.75 | | 25.86 | 24.48 | | |
| 232 | T | - | 62.9 | 176.691 | 70.789 | 26.91 | | | | | |
| 233 | D | - | - | - | - | - | | | | | |
| 234 | Q | - | - | - | - | - | | - | | | |
| 235 | G | 108.8 | 46.7 | 173.726 | | | | | | | |
| 236 | I | - | 60.2 | 177.1 | 36.59 | 31.07 | - | - | | | |
| 237 | A | - | - | - | - | | | | | | |

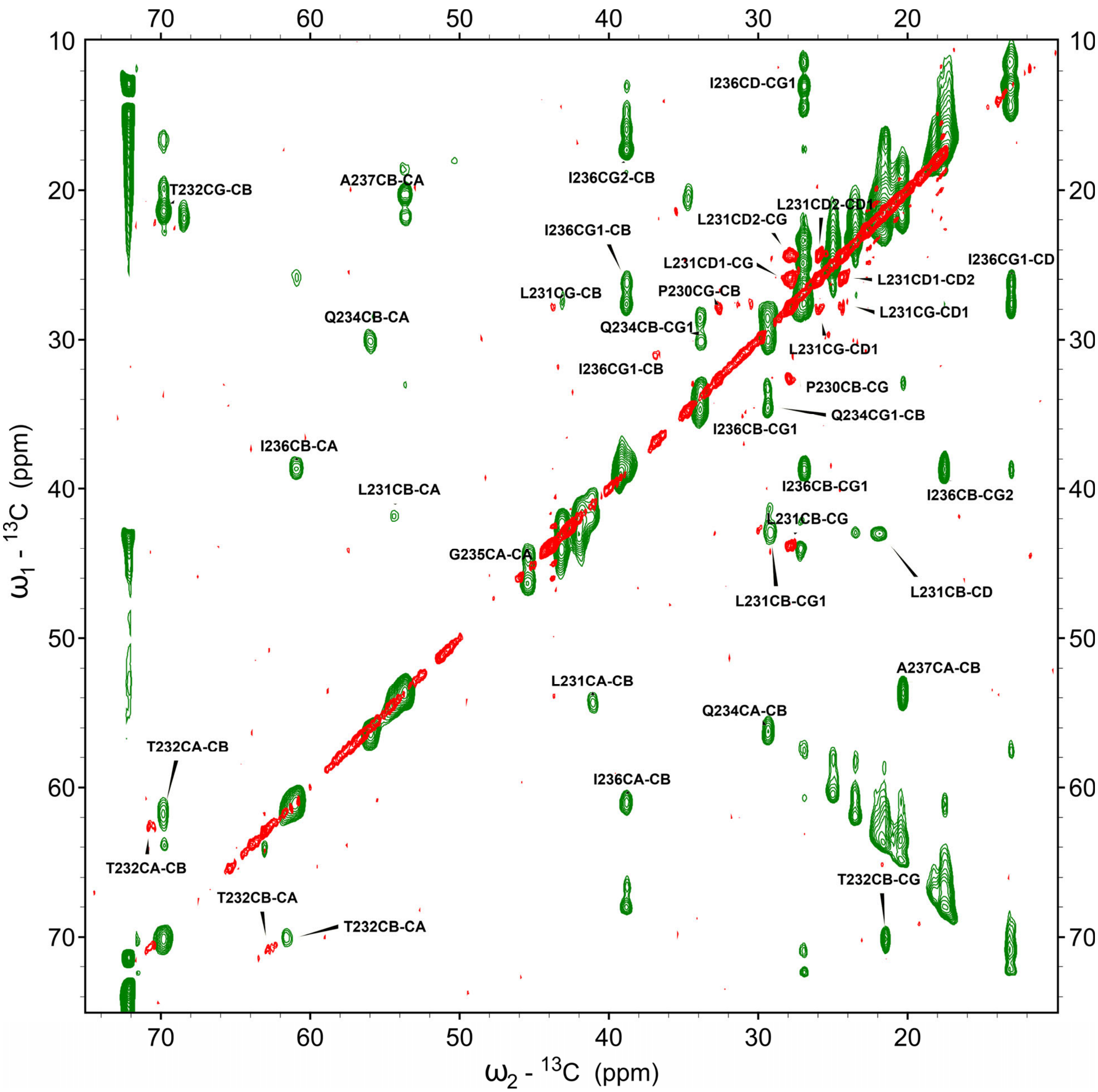


Figure S1. Several residue chemical shifts were identified by INEPT-TOBSY experiments. Overlay of 2D $^{13}C$-$^{13}C$ spectra taken from tubular (green) and T=1 capsid (red) assembly samples.

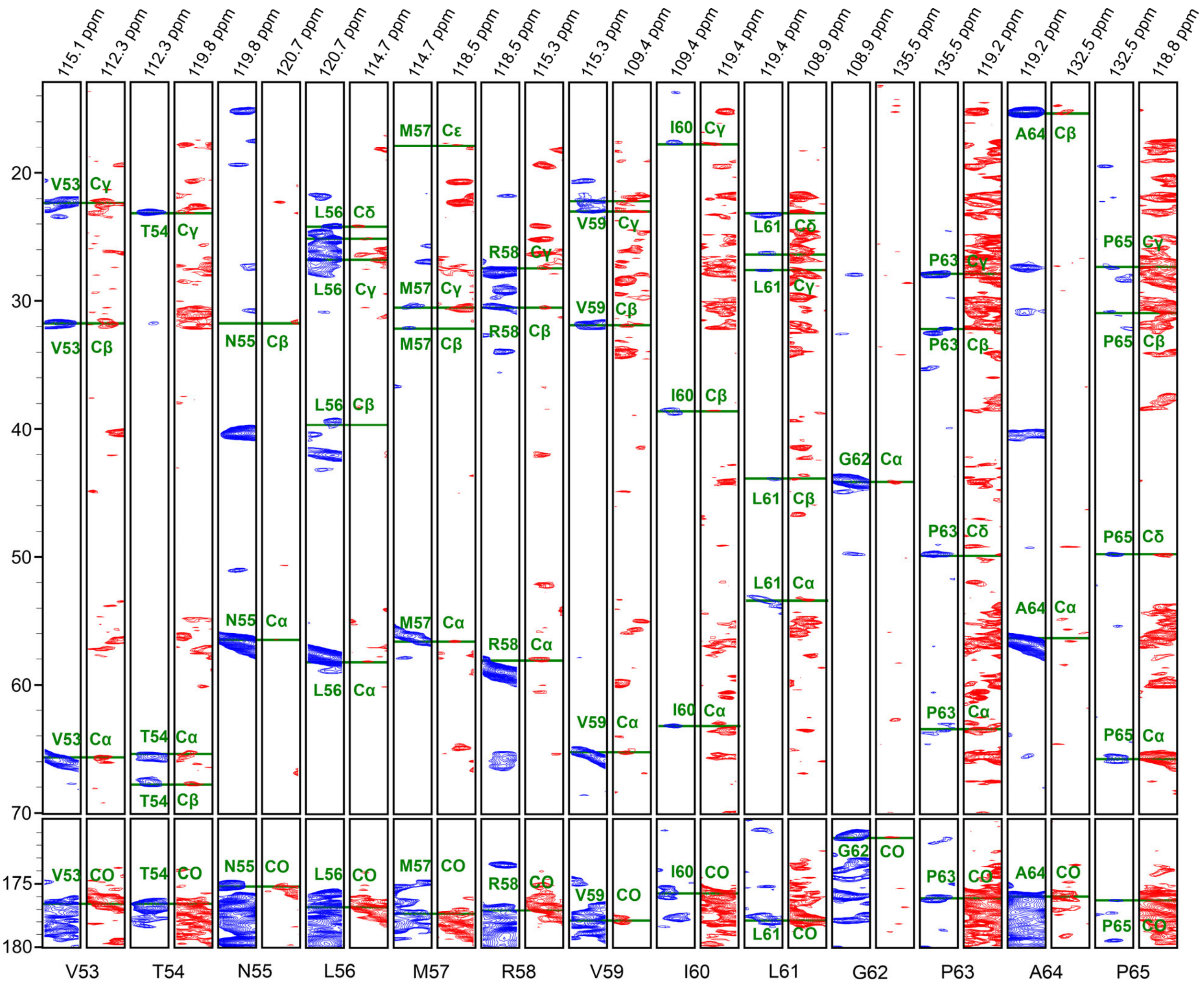


Figure S2. Sequential assignments for residues V53 to P65 in the T=1 capsid assembly illustrated by the backbone walk technique, connecting slices of the 3D NCACX (blue) and NCOCX (red).

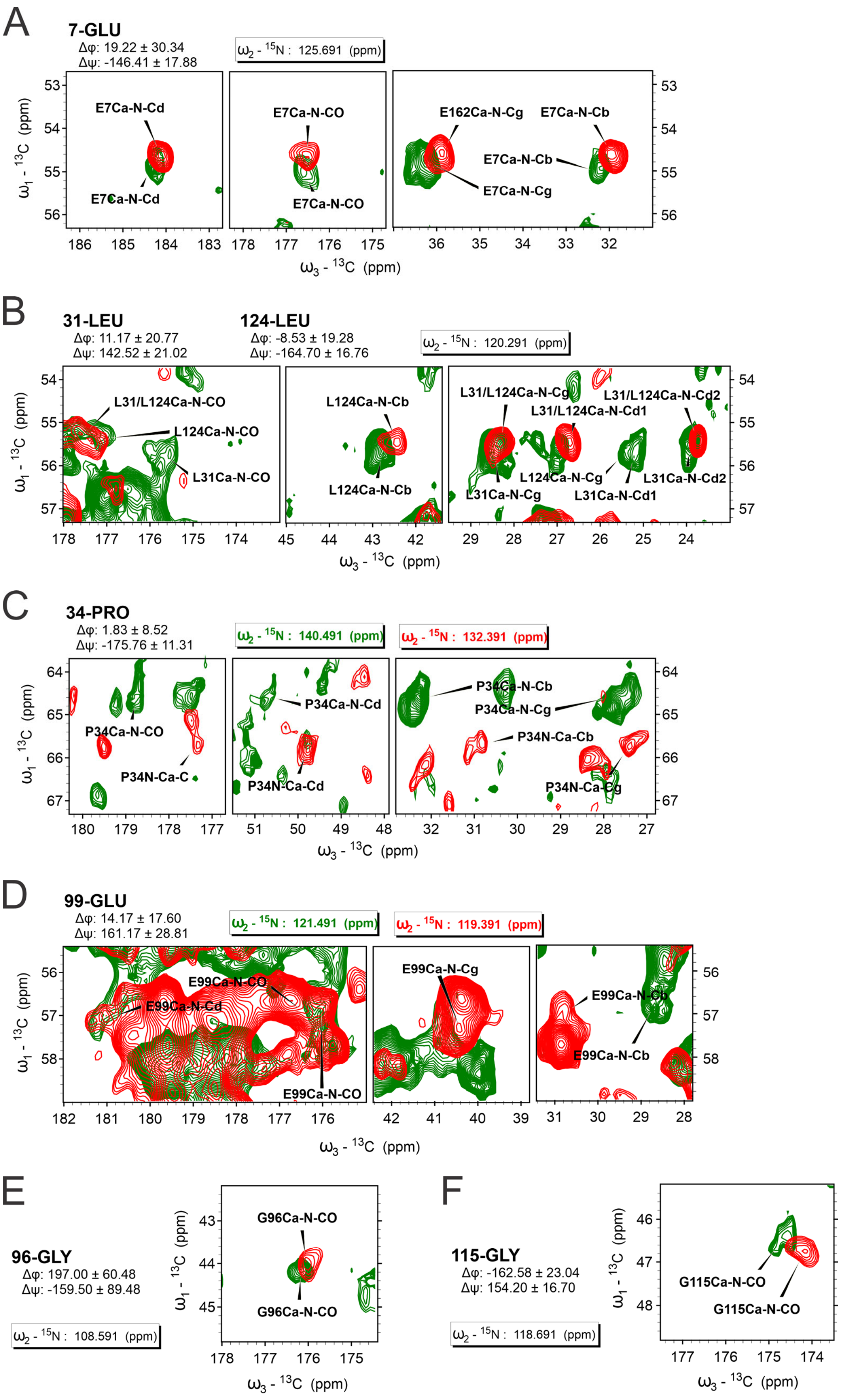

A
7-GLU
Δφ: 19.22 ± 30.34
Δψ: -146.41 ± 17.88
ω2 - 15N : 125.691 (ppm)
E7Ca-N-Cd
E7Ca-N-Cd
E7Ca-N-CO
E7Ca-N-CO
E162Ca-N-Cg
E7Ca-N-Cb
E7Ca-N-Cb
E7Ca-N-Cg
ω1 - 13C (ppm)
ω3 - 13C (ppm)
B
31-LEU
Δφ: 11.17 ± 20.77
Δψ: 142.52 ± 21.02
124-LEU
Δφ: -8.53 ± 19.28
Δψ: -164.70 ± 16.76
ω2 - 15N : 120.291 (ppm)
L31/L124Ca-N-CO
L124Ca-N-CO
L31Ca-N-CO
L124Ca-N-Cb
L124Ca-N-Cb
L31/L124Ca-N-Cg
L31/L124Ca-N-Cd2
L31/L124Ca-N-Cd1
L124Ca-N-Cg
L31Ca-N-Cd2
L31Ca-N-Cg
L31Ca-N-Cd1
ω1 - 13C (ppm)
ω3 - 13C (ppm)
C
34-PRO
Δφ: 1.83 ± 8.52
Δψ: -175.76 ± 11.31
ω2 - 15N : 140.491 (ppm)
ω2 - 15N : 132.391 (ppm)
P34Ca-N-CO
P34N-Ca-C
P34Ca-N-Cd
P34N-Ca-Cd
P34Ca-N-Cb
P34Ca-N-Cg
P34N-Ca-Cb
P34N-Ca-Cg
ω1 - 13C (ppm)
ω3 - 13C (ppm)
D
99-GLU
Δφ: 14.17 ± 17.60
Δψ: 161.17 ± 28.81
ω2 - 15N : 121.491 (ppm)
ω2 - 15N : 119.391 (ppm)
E99Ca-N-CO
E99Ca-N-Cd
E99Ca-N-CO
E99Ca-N-Cg
E99Ca-N-Cb
E99Ca-N-Cb
ω1 - 13C (ppm)
ω3 - 13C (ppm)
E
96-GLY
Δφ: 197.00 ± 60.48
Δψ: -159.50 ± 89.48
ω2 - 15N : 108.591 (ppm)
G96Ca-N-CO
G96Ca-N-CO
ω1 - 13C (ppm)
ω3 - 13C (ppm)
F
115-GLY
Δφ: -162.58 ± 23.04
Δψ: 154.20 ± 16.70
ω2 - 15N : 118.691 (ppm)
G115Ca-N-CO
G115Ca-N-CO
ω1 - 13C (ppm)
ω3 - 13C (ppm)

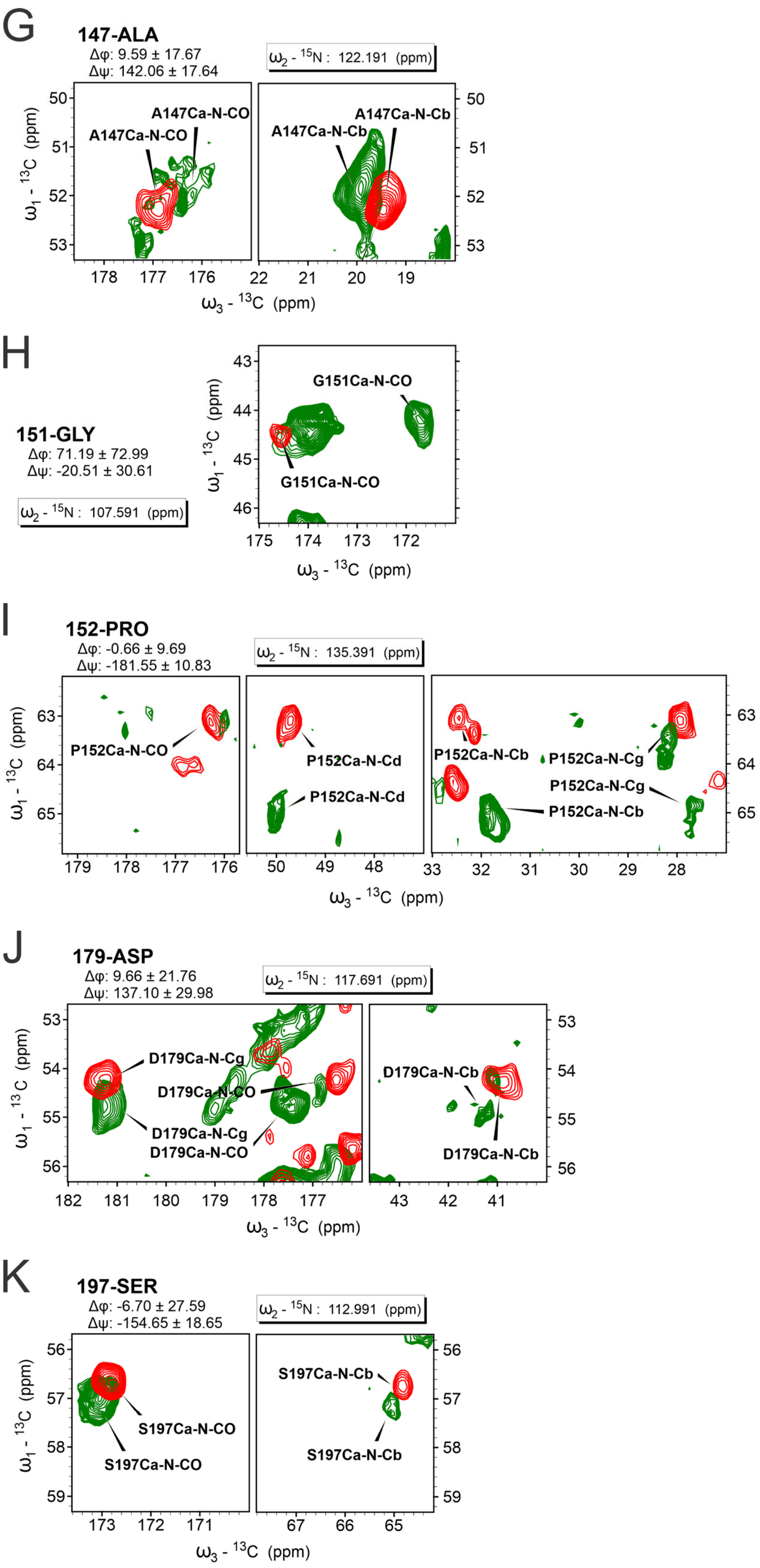
G
147-ALA
Δφ: 9.59 ± 17.67
Δψ: 142.06 ± 17.64
ω2 - 15N : 122.191 (ppm)
A147Ca-N-CO
A147Ca-N-CO
A147Ca-N-Cb
A147Ca-N-Cb
ω1 - 13C (ppm)
ω3 - 13C (ppm)
H
151-GLY
Δφ: 71.19 ± 72.99
Δψ: -20.51 ± 30.61
ω2 - 15N : 107.591 (ppm)
G151Ca-N-CO
G151Ca-N-CO
ω1 - 13C (ppm)
ω3 - 13C (ppm)
I
152-PRO
Δφ: -0.66 ± 9.69
Δψ: -181.55 ± 10.83
ω2 - 15N : 135.391 (ppm)
P152Ca-N-CO
P152Ca-N-Cd
P152Ca-N-Cd
P152Ca-N-Cb
P152Ca-N-Cg
P152Ca-N-Cg
P152Ca-N-Cb
ω1 - 13C (ppm)
ω3 - 13C (ppm)
J
179-ASP
Δφ: 9.66 ± 21.76
Δψ: 137.10 ± 29.98
ω2 - 15N : 117.691 (ppm)
D179Ca-N-Cg
D179Ca-N-CO
D179Ca-N-Cg
D179Ca-N-CO
D179Ca-N-Cb
D179Ca-N-Cb
ω1 - 13C (ppm)
ω3 - 13C (ppm)
K
197-SER
Δφ: -6.70 ± 27.59
Δψ: -154.65 ± 18.65
ω2 - 15N : 112.991 (ppm)
S197Ca-N-CO
S197Ca-N-CO
S197Ca-N-Cb
S197Ca-N-Cb
ω1 - 13C (ppm)
ω3 - 13C (ppm)

Figure S3. (A-K) Spectral slices of residues that exhibit large difference in torsion angle between T=1 capsid and tubular assemblies. Contour plot overlay taken from $^{15}N$ plane slices of 3D NCACX spectra, with red contour corresponding to the T=1 capsid assembly and green contour corresponding to the tubular assembly. Residues Pro50 and Ala150 were only assigned in tubular and T=1 capsid assignments respectively and thus do not have an overlay shown. Torsion angle differences (T=1 capsid minus tubular) provided.

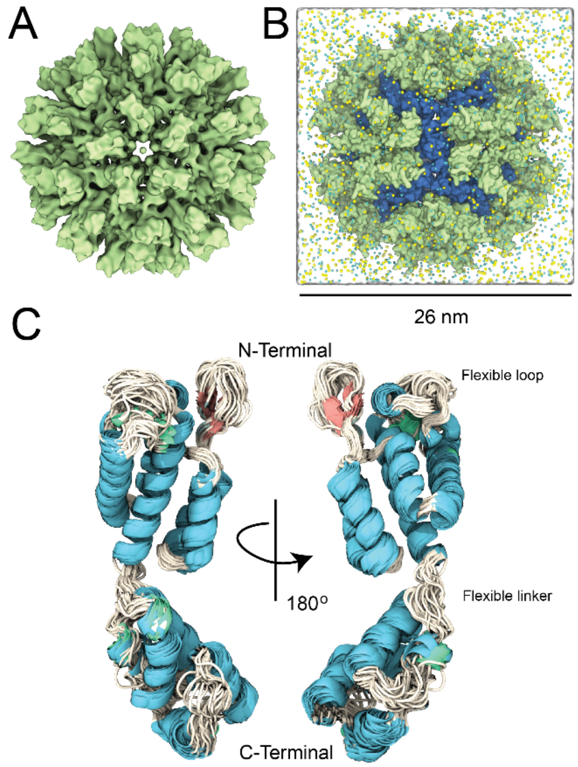


Figure S4. Structural model of the RSV CA T=1 capsid, obtained by simulations to combine the 8.5 A electron density map (EMDB 5772) with our ssNMR results. (B). The T=1 system comprising 60 monomeric RSV CA with secondary structure defined by our NMR results, solvated in 150 mM salt. Each monomer comprises 226 residues of RSV CA, with residues from the disordered tail excluded in the model. (C) Alignments of the 60 monomers in the T=1 capsid by their backbone atoms.

### References:

[1] Bowzard, J. B., Wills, J. W., and Craven, R. C. (2001) Second-site suppressors of Rous sarcoma virus CA mutations: Evidence for interdomain interactions, *Journal of Virology 75*, 6850-6856.

[2] Jeon, J., Qiao, X., Hung, I., Mitra, A. K., Desfosses, A., Huang, D., Gor'kov, P. L., Craven, R. C., Kingston, R. L., Gan, Z. H., Zhu, F. Q., and Chen, B. (2017) Structural Model of the Tubular Assembly of the Rous Sarcoma Virus Capsid Protein, *Journal of the American Chemical Society 139*, 2006-2013.

[3] Delaglio, F., Grzesiek, S., Vuister, G. W., Zhu, G., Pfeifer, J., and Bax, A. (1995) NMRPIPE - A MULTIDIMENSIONAL SPECTRAL PROCESSING SYSTEM BASED ON UNIX PIPES, *Journal of Biomolecular Nmr 6*, 277-293.

[4] Lee, W., Tonelli, M., and Markley, J. L. (2015) NMRFAM-SPARKY: enhanced software for biomolecular NMR spectroscopy, *Bioinformatics 31*, 1325-1327.

[5] Phillips, J. C., Braun, R., Wang, W., Gumbart, J., Tajkhorshid, E., Villa, E., Chipot, C., Skeel, R. D., Kale, L., and Schulten, K. (2005) Scalable molecular dynamics with NAMD, *Journal of computational chemistry 26*, 1781-1802.

[6] Huang, J., Rauscher, S., Nawrocki, G., Ran, T., Feig, M., de Groot, B. L., Grubmüller, H., and MacKerell Jr, A. D. (2017) CHARMM36m: an improved force field for folded and intrinsically disordered proteins, *Nature methods 14*, 71.

[7] Shen, Y., and Bax, A. (2015) Protein structural information derived from NMR chemical shift with the neural network program TALOS-N, In *Artificial neural networks*, pp 17-32, Springer.

[8] Trabuco, L. G., Villa, E., Mitra, K., Frank, J., and Schulten, K. (2008) Flexible fitting of atomic structures into electron microscopy maps using molecular dynamics, *Structure 16*, 673-683.

[9] Søndergaard, C. R., Olsson, M. H. M., Rostkowski, M., and Jensen, J. H. (2011) Improved Treatment of Ligands and Coupling Effects in Empirical Calculation and Rationalization of pKa Values, *Journal of Chemical Theory and Computation 7*, 2284-2295.

[10] Keller, P. W., Huang, R. K., England, M. R., Waki, K., Cheng, N. Q., Heymann, J. B., Craven, R. C., Freed, E. O., and Steven, A. C. (2013) A Two-Pronged Structural Analysis of Retroviral Maturation Indicates that Core Formation Proceeds by a Disassembly-Reassembly Pathway Rather than a Displacive Transition, *Journal of Virology 87*, 13655-13664.

[11] Pettersen, E. F., Goddard, T. D., Huang, C. C., Couch, G. S., Greenblatt, D. M., Meng, E. C., and Ferrin, T. E. (2004) UCSF chimera - A visualization system for exploratory research and analysis, *Journal of Computational Chemistry 25*, 1605-1612.

[12] Humphrey, W., Dalke, A., and Schulten, K. (1996) VMD: visual molecular dynamics, *Journal of molecular graphics 14*, 33-38.

[13] Darden, T., York, D., and Pedersen, L. (1993) Particle mesh Ewald: An N· log (N) method for Ewald sums in large systems, *The Journal of chemical physics 98*, 10089-10092.

[14] Essmann, U., Perera, L., Berkowitz, M. L., Darden, T., Lee, H., and Pedersen, L. G. (1995) A smooth particle mesh Ewald method, *The Journal of chemical physics 103*, 8577-8593.

[15] Skeel, R. D., and Biesiadecki, J. J. (1994) Symplectic integration with variable stepsize, *Annals of Numerical Mathematics 1*, 191-198.

[16] Bussi, G., Donadio, D., and Parrinello, M. (2007) Canonical sampling through velocity rescaling, *The Journal of chemical physics 126*, 014101.

[17] Kräutler, V., Van Gunsteren, W. F., and Hünenberger, P. H. (2001) A fast SHAKE algorithm to solve distance constraint equations for small molecules in molecular dynamics simulations, *Journal of computational chemistry 22*, 501-508.

[18] Andersen, H. C. (1983) Rattle: A "velocity" version of the shake algorithm for molecular dynamics calculations, *Journal of Computational Physics 52*, 24-34.
